# Human neuronal networks on micro-electrode arrays are a highly robust tool to study disease-specific genotype-phenotype correlations *in vitro*

**DOI:** 10.1101/2021.01.20.427439

**Authors:** B. Mossink, A.H.A. Verboven, E.J.H. van Hugte, T.M. Klein Gunnewiek, G. Parodi, K. Linda, C. Schoenmaker, T. Kleefstra, T. Kozicz, H. van Bokhoven, D. Schubert, N. Nadif Kasri, M. Frega

## Abstract

Micro-electrode arrays (MEAs) are increasingly used to characterize neuronal network activity of human induced pluripotent stem-cell (hiPSC)-derived neurons. Despite their gain in popularity, MEA recordings from hiPSC-derived neuronal networks are not always used to their full potential in respect to experimental design, execution and data analysis. Therefore, we benchmarked the robustness and sensitivity of MEA-derived neuronal activity patterns derived from ten healthy individual control lines. We provide recommendations on experimental design and analysis to achieve standardization. With such standardization, MEAs can be used as a reliable platform to distinguish (disease-specific) network phenotypes. In conclusion, we show that MEAs are a powerful and robust tool to uncover functional neuronal network phenotypes from hiPSC-derived neuronal networks, and provide an important resource to advance the hiPSC field towards the use of MEAs for disease-phenotyping and drug discovery.

## Introduction

*In vitro* neuronal models have become an important tool for studying the complex communication of healthy and diseased neuronal circuits. In particular, the possibility to observe, measure and manipulate the electrical activity exhibited by neuronal populations can give insights into neuronal network development and organization (Novellino et al. 2011). Micro-electrode arrays (MEAs) are cell culture dishes with embedded micro-electrodes that allow non-invasive measurement of neuronal network activity. MEAs have been extensively used to measure activity from a range of different neuronal culture systems, for example primary cell cultures, brain slices or intact retinas, mainly from rodent origin (McConnell et al. 2012). With the advancements in human induced pluripotent stem cell (hiPSC) technology, the differentiation of human neurons from somatic cells became possible, allowing neuronal network phenotyping with a model system that more closely resembles the human brain. HiPSC-derived neuronal networks on MEA mimic the activity pattern of rodent neuronal networks, including a stable state of synchronized network bursting, suggesting that they successfully develop into functional neuronal networks (Odawara et al. 2014, Fukushima et al. 2016, Odawara et al. 2016, Kayama et al. 2018, Frega et al. 2019, Sasaki et al. 2019). In addition, the development of comprehensive MEA analysis software has increased over the years, simplifying the extraction of parameters that describe the pattern of neuronal activity observed in MEA recordings. These advancements in both human neuronal culturing systems and MEA analysis software contributed to the popularity of MEA technology to study neuronal network phenotypes in health and disease (Wainger et al. 2014, Deneault et al. 2019, Frega et al. 2019, Gunnewiek et al. 2020).

Despite their increasing popularity, MEA measurements are not always used to their full potential to investigate hiPSC-derived neuronal network characteristics. HiPSC-derived neuronal networks have not been benchmarked as extensively as that of rodent neuronal cultures. Because of the lack of standardization, little is currently known about how changes in cell-culture conditions influence batch-to-batch consistency, and how different hiPSC-derived neuronal lines affect reproducibility (Engle et al. 2018). It was previously advised to use multiple lines or isogenic sets to reliably determine a disease phenotype, since differences in genetic background between hiPSCs donors dominate the variance at the transcriptional level (Germain et al. 2017). However, little is known about the amount of cell lines needed to distinguish a phenotype on MEAs, or about the effect of genetic background on hiPSC-derived neuronal network function. In addition to experimental design, data analysis remains a hurdle, even though the extraction of different MEA parameters became easier. Studies often quantify the general neuronal network activity by a single parameter (i.e. mean firing rate), thereby failing to explain the complex network characteristics. Finally, cell culture practices are not always optimized and thus mature networks, showing network synchronicity, are not always obtained. In summary, the question remains how reproducible and comparable MEA recordings are within and between different lines, different researchers, and across different batches or developmental time points, illustrating the need for a quality standard.

In this study we provide a set of recommendations for the design, analysis, and interpretation of hiPSC-derived neuronal studies on MEAs. We performed a meta-analysis of MEA recordings from excitatory neuronal networks derived from hiPSCs through one of the most widely used differentiation protocols (i.e. *Ngn2* induction (Zhang et al. 2013, Frega et al. 2017)). Specifically, we used hiPSCs derived from ten healthy subjects, which were cultured by different researchers over a period of several years. We show that control neuronal networks cultured on MEAs are highly reproducible between different recordings and control lines, and identify the most robust MEA parameters to describe neuronal network activity and organization. When pooling data from all control lines together, the functional activity of control neuronal networks is not largely influenced by biological differences between donors (i.e. age, sex). Using neuronal networks affected by genetic aberrations causing Kleefstra syndrome (KS) or Mitochondrial encephalopathy, lactic acidosis, and stroke-like episodes (MELAS), we show that the MEA platform is a powerful tool to identify genotype-phenotype correlations.

## Results

### Excitatory neurons derived from healthy subjects show a comparable phenotype on MEA

To investigate if network activity from hiPSC-derived *Ngn2*-induced excitatory neurons are comparable, we performed a meta-analysis on MEA data derived from multiple control lines used in our lab (Frega et al. 2019, Gunnewiek et al. 2020, Mossink et al. 2020). We generated hiPSCs from fibroblast skin biopsies from ten individuals, five males and five females, with a mean age of 33.5 years (**Fig. 1a**). We extracted in total 17 MEA parameters to describe the neuronal network activity and connectivity (**Table S1**).

**Figure 1.**
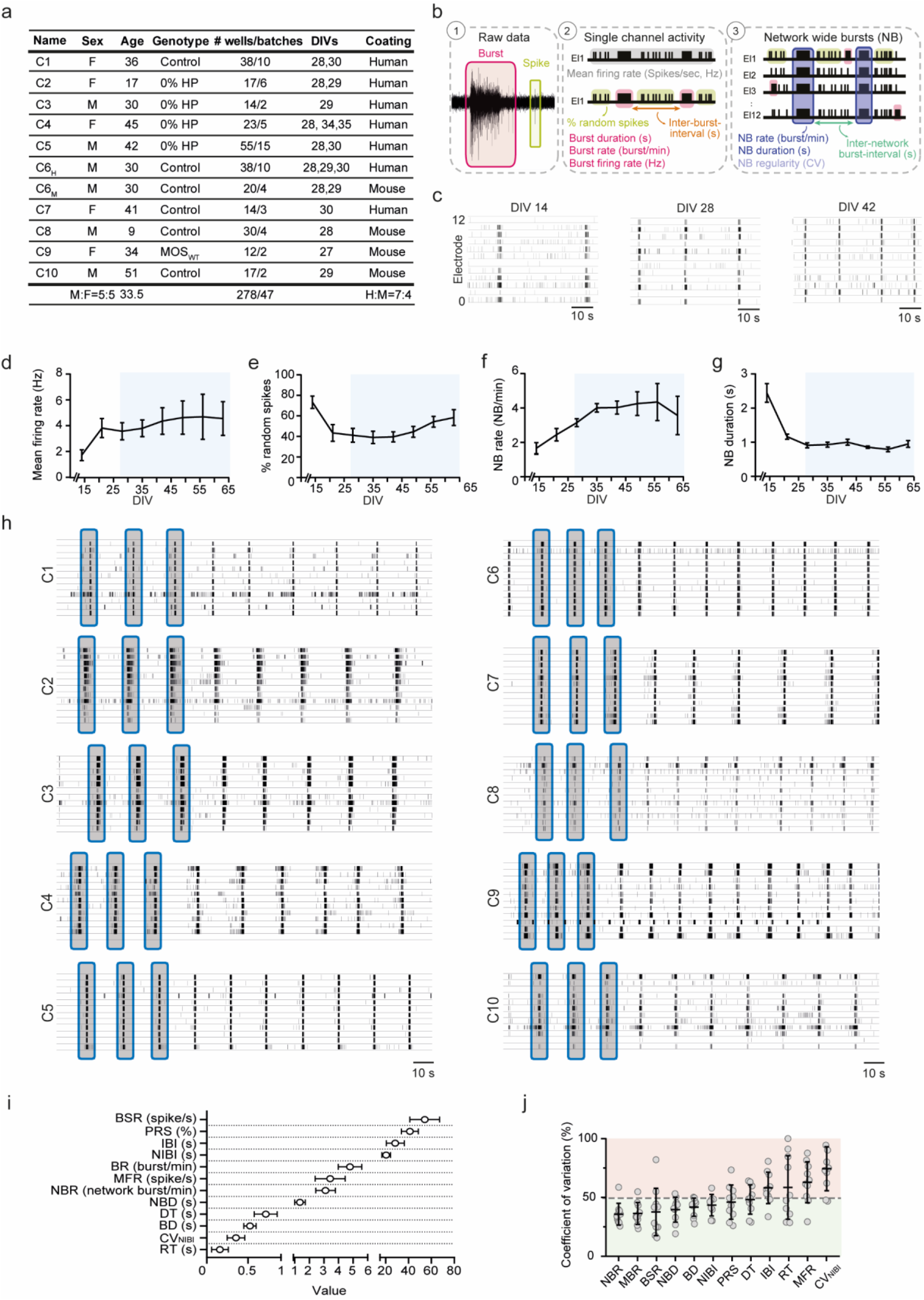
Control neuronal networks show a stable phenotype on MEA. (**a**) Information regarding the 10 control lines used in this study. C_6_ was recorded on two substrates (H = human laminin and M = mouse laminin). Number of wells represent total number of wells recorded for that line between DIV27-35, including the number of batches. Some batches overlap between lines. (**b**) Schematic overview of extracted parameters from MEA (see Table S1). (**c**) Representative raster plots of line C_6_ showing 60 seconds of electrophysiological activity across development (DIV14-42). (**d-g**) Neuronal network parameters (of line C_6_) develop to reach a certain plateau after DIV 27 (blue box) for (**d**) MFR, (**e**) PRS, (**f**) NBR and (**g**) NBD. (**h**) Representative raster plots of 10 control lines showing 3 minutes of electrophysiological activity on MEAs (**i**) Graph showing the range in which MEA parameters of all 10 control lines behave (mean ± 95% confidence interval). Values are first averaged per control line, and then averaged across all control lines. (**j**) Percent coefficient of variation explaining the stability of the respective MEA parameter across all 10 control lines (mean ± standard deviation of the mean). N = 278 wells (**Table S2**). DIV = days in vitro, MFR = mean firing rate, PRS = percentage of random spikes, BR = mean burst rate, BD = mean burst duration, BSR = Burst spike rate, IBI = inter-burst interval, NBR = network burst rate, NBD = network burst duration, NIBI = Network burst IBI, CV_NIBI_ = coefficient of variation of all NIBI’s representing the regularity of the NB, RT = Rise time, DT = decay time. All means are reported in supplementary table 2.

During the first two weeks of differentiation, neuronal network activity on MEAs primarily consisted of random spikes (single action potentials) and bursts (high frequency action potentials), which during development organized into network bursts (rhythmic, synchronous events) (**Fig. 1b, c**). During maturation, *Ngn2*-induced neuronal networks displayed an increase of firing and (network) bursting activities, and a decrease in (network) burst duration and random spike activity (**Fig. 1d-g, S1a-e**). From 27 days in vitro (DIV) onwards, these parameters plateaued, and neuronal network activity remained stable (blue boxes). Because these neuronal networks were generally measured between DIV 27-35, and network activity remained stable after DIV 27, we pooled data in this developmental window (n_wells_ = 278).

At this specific time point, we observed similar patterns of activity and connectivity across all control lines (**Fig. 1h, Fig. S1f-o, Fig. S2, Fig. S3**). We determined the specific range of values for each MEA parameter describing neuronal network characteristics using data from all control lines. The neuronal network functioning of control lines was within a tightly controlled range (**Fig. 1i, Fig. S3f, Table S2**). Control neuronal networks showed a general level of activity of 3.5 ± 0.2 spike/s and synchronous events appeared with a frequency of 3.2 ± 0.1 network bursts/min and a duration of 1.28 ± 0.04 s. We did observe slight differences between individual control lines for some parameters. For example, at similar development ages, control lines C_2_ and C_9_ exhibited synchronous events at different frequencies as compared to the other controls (i.e. 1.4 ± 0.2 and 4.6 ± 0.3 network bursts/min for C_2_ and C_9_, respectively, **Fig. S1m**). This stresses the need of using multiple lines to uncover the full phenotypic spectrum of control neuronal networks. Taken together, these results indicate that neuronal networks on MEA show similar patterns of activity across multiple control lines.

Next, we investigated the variability of the MEA parameters within our control dataset to identify the most robust parameter(s) (i.e. coefficient of variation lower than 50% as cutoff, **Fig. 1j, Fig. S3g**). Our results indicated that certain parameters were more stable (i.e. frequency and duration of the (network) bursts, (N)BR and (N)BD respectively), whereas others were more variable (i.e. mean firing rate (MFR), regularity of the network burst appearance (CV_NIBI_), C_0_, link weight). In most of the hiPSC-based MEA studies the MFR has been used as the main and only parameter, which may confound the characterization and interpretation of the neuronal network behavior. Besides the fact that MFR is one of the most variable parameters we analyzed, it is highly dependent on cell density (Biffi et al. 2013) and it only describes the general level of activity, without any information about network organization (i.e. synchronization, connectivity). Multiple MEA parameters describing both general activity and bursting behavior should be included to obtain a comprehensive characterization of neuronal network behavior.

### Confounding factors in experimental design, culturing and analysis that influence the reliability of neuronal network recordings

When we combined all MEA parameters in a principal component analysis (PCA), we did not observe clear clustering based on hiPSC line (**Fig. 2a**). This indicated that there was no consistent line-specific difference at the functional level. To guarantee these reliable neuronal network recordings, we explored which confounding factors can introduce or increase variation. Our analyses of the datasets showed that sex and age of the original fibroblast donor had no major effect on the neuronal network phenotype variability (**Fig. 2b, c**). Furthermore, we found no clear clustering based on DIV when the cultures reached a stable developmental stage (i.e. DIV 27-35, **Fig. 2d**). However, neuronal networks measured on earlier time points (DIV 14-24) clustered away from measurements performed after DIV 28 (**Fig. S1p**). Thus, pooling data from different developmental stages should be avoided since it likely introduces variation in the data.

**Figure 2.**
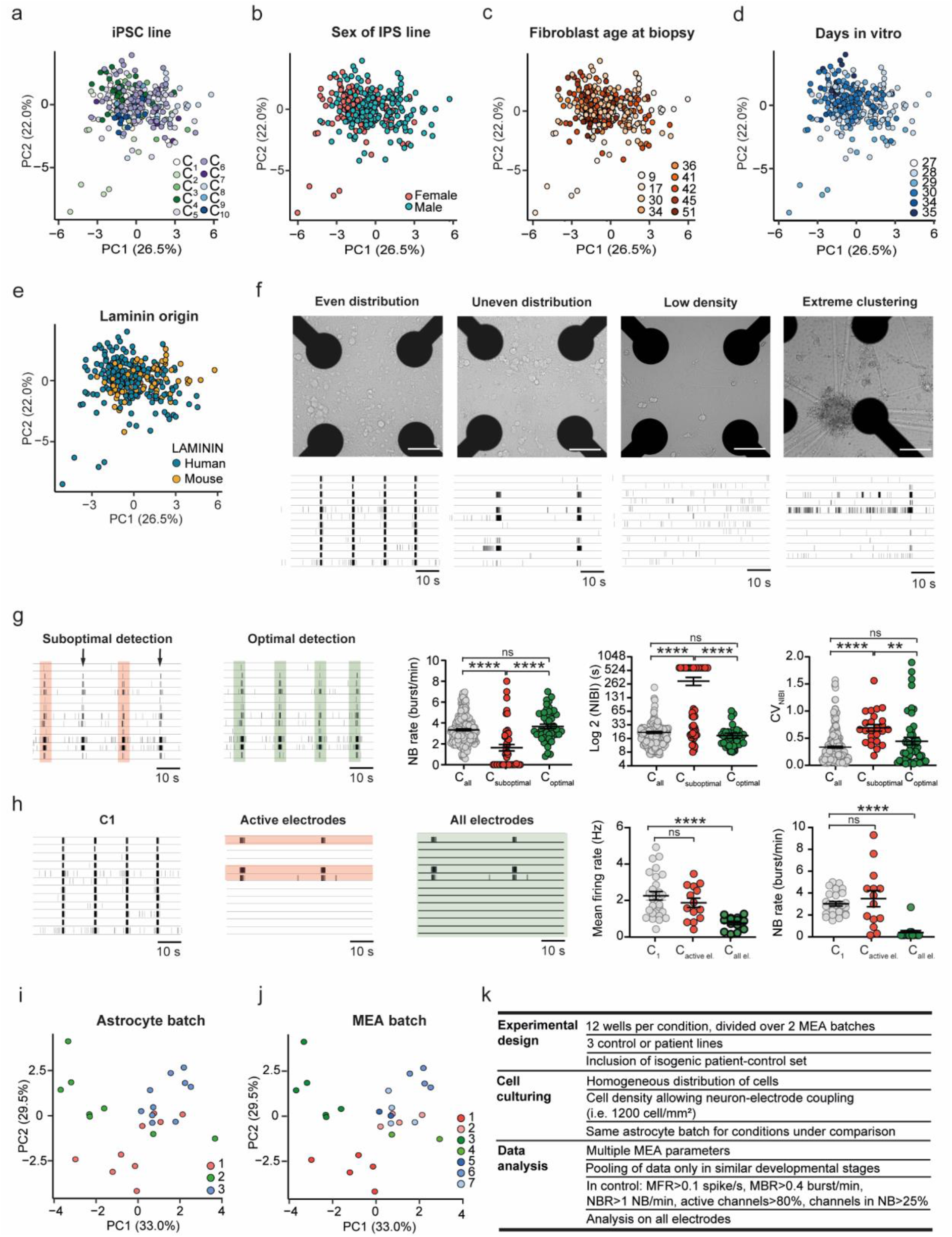
Variables that influence neuronal network phenotype. (**a-e**) PCA plot on all parameters showing data of all control lines pooled from DIV27-35 (**a**) color-coded by line, (**b**) color-coded by sex, (**c**) color-coded by the fibroblast age at biopsy, (**d**) color-coded by DIV, and (**e**) color-coded by laminin origin. (**f**) Representative images of neuronal cultures grown at different densities and distributions (even, uneven and low densities, and extreme clustering) and representative raster plots showing 1 minute of activity exhibited by neuronal networks in each condition. (**g**) Representative raster plots of a well in which the network burst detection was adapted to detect all network burst present. Colored bars represent the detected network burst by software. Comparison of the MEA parameters NBR, NIBI (on log2 scale) and CV_NIBI_ between control pool (C_all_ grey), wells in which not all network burst have been detected (C_suboptimal_, red) and the same wells when optimal detection have been performed (C_optimal_, green). Kruskal Wallis Anova with Dunn’s correction for multiple testing was used to compare between control lines. (**h**) Representative raster plots of C1 and a well in which only a few channels are active. In the figure, the electrodes used for analysis are highlighted (green and red for 3 active electrodes and all electrodes, respectively). Comparison of the MEA parameters MFR and NBR between C1, well in which the analysis has been performed only on active electrode (C_active el,_ red) and the same well when the analysis has been performed on all electrodes (C_all el,_ green). One-way Anova with Tukey correction for multiple testing was used to compare between control lines. (**i-j**) PCA plot on all parameters showing data of (**i**) one control line (C1) color-coded by astrocyte batch and (**j**) one control line (C1) and color-coded by MEA batch. (**k**) List of recommendations about experimental design, cell culturing and data analysis. *p* = 0.05*, *p* = 0.01 ** and *p* = 0.001 ***. DIV = days in vitro, MFR = mean firing rate, PRS = percentage of random spikes, NBR = network burst rate, NBD = network burst duration, NIBI = Network burst IBI, CV_NIBI_ = coefficient of variation of all NIBI’s representing the regularity of the NB. All means, *p*-values and used statistic tests are reported in supplementary Table S4.

Next, we explored whether culturing conditions can be a confounding factor on neuronal network behavior. First, we observed no clear difference between neuronal networks grown on two types of coatings used for cell culturing (mouse or human laminin) at the stable developmental stage (**Fig. 2e**). However, different developmental trajectories have been observed in neuronal networks grown on mouse and human laminin (Pujadas et al. 2012, Hyysalo et al. 2017), thus pooling and comparison of data from different coatings can affect their comparability. Another culturing variable that could influence network activity is cell distribution. With low resolution MEA systems (i.e. 12 electrodes spaced 300 μm apart) the activity recorded from the electrodes originates from multiple neurons. Therefore, homogeneous distribution of cells on each electrode should be achieved to reduce variability. Indeed, we found that changes in cell density and distribution affected neuronal network functionality (**Fig. 2f**). Whereas an even distribution of neurons on all electrodes was accompanied by synchronous activity involving all channels, an uneven distribution can lead to events involving only a few channels (**Fig. 2f**, second panel). In addition, neuronal networks with (extreme) low densities exhibited less frequent events (**Figure 2f**, third panel) or only random spikes, since too few neurons were grown on the electrode (**Fig. 2f**, third panel). Cell clustering led to highly frequent local activity recorded only detected by the electrodes close to the cluster (**Fig. 2f**, fourth panel). Thus, cell density and distribution should be consistent between neuronal networks to achieve a comparable network pattern. A final cell density allowing a proper neuron-electrode coupling should be chosen (e.g. 1200 cells/mm^2^ was used while generating this dataset). Neuronal networks with low cell density or uneven distribution of cells should be excluded from the analysis.

In addition to culturing conditions, we found that accurate data analysis depended on the selection of proper analysis settings. Suboptimal network burst detection influenced several parameters, and consequently resulted in a faulty quantification of the neuronal network organization, while the raster plot was similar to all the other controls (**Fig. 2g**, C_suboptimal_ vs C_all_). When an optimal detection was performed on the same dataset, no difference between the two groups was present (C_optimal_ vs C_all_), as expected from the raster plot. Similarly, data analysis performed on individual active electrodes lead to erroneous results when culturing conditions were not optimal. When we analyzed only the active electrodes in wells with uneven densities, we obtained similar outcomes in activity patterns as in wells with optimal density in which all electrodes were analyzed, resulting in an incorrect representation of the actual neuronal network (**Fig. 2h**, C_active el._ vs C_1_). Analysis on all electrodes however provided a correct image of the neuronal network phenotype (C_all el._ vs C_1_). Thus, stringent criteria should be used when performing data analysis. Control neuronal networks should display at least certain levels of activity to be included in a further analysis, including a MFR > 0.1 spike/s, a burst rate > 0.4 bursts/min and a network burst rate > 1 network burst/min and synchronous activity should be observed in most of the channels. General activity (i.e. spikes) should be detected in at least 80 % of the electrodes and analysis should be performed on all electrodes rather than only on the active ones.

Finally, we investigated the effect of both independent astrocyte batches and MEA batches (i.e. independent neuronal preparations on MEA) on the neuronal network behavior. PCA showed samples cluster based on astrocyte batch (**Fig. 2i**), indicating that astrocyte batch affected the neuronal network phenotype. We also observed clustering of samples based on the MEA batch, illustrating the effect of individual neuronal preparations on the neuronal network phenotype, independent of the astrocyte batch used (**Fig. 2j**). These results stress the need for using multiple experimental batches to correct for this technical variation (i.e. at least two MEA batches with astrocytes belonging to the same batch).

In summary, our results indicate that certain standards should be followed to ensure that reliable data is obtained from MEA experiments (**Fig. 2k**). To generate reproducible neuronal control network phenotypes, one needs to (i) culture sufficient neurons that are homogenously distributed, (ii) properly select the detection settings, (iii) pool data only in a certain developmental time windows, and (iv) use sufficient experimental batches.

### The MEA system is a reliable platform for disease phenotyping

To confirm that control neuronal networks can be used as a platform to perform disease phenotyping, we compared patient neuronal network activity from two neurodevelopmental disorders (NDD) with the total control pool. In particular, we re-analyzed our previously published data from three patients with MELAS syndrome (two females and one male, mean age of 34.7 years) (Gunnewiek et al. 2020) and four patients with Kleefstra syndrome (KS) (three females and one male, mean age of 27.5 years) (Frega, Linda et al. 2019) (**Fig. 3a**). Since control neuronal networks were stable between DIV 27-35, recordings from MELAS patient lines (n_wells_ = 112), as well as KS patient lines (n_wells_ = 58), were pooled in the same time window.

**Figure 3.**
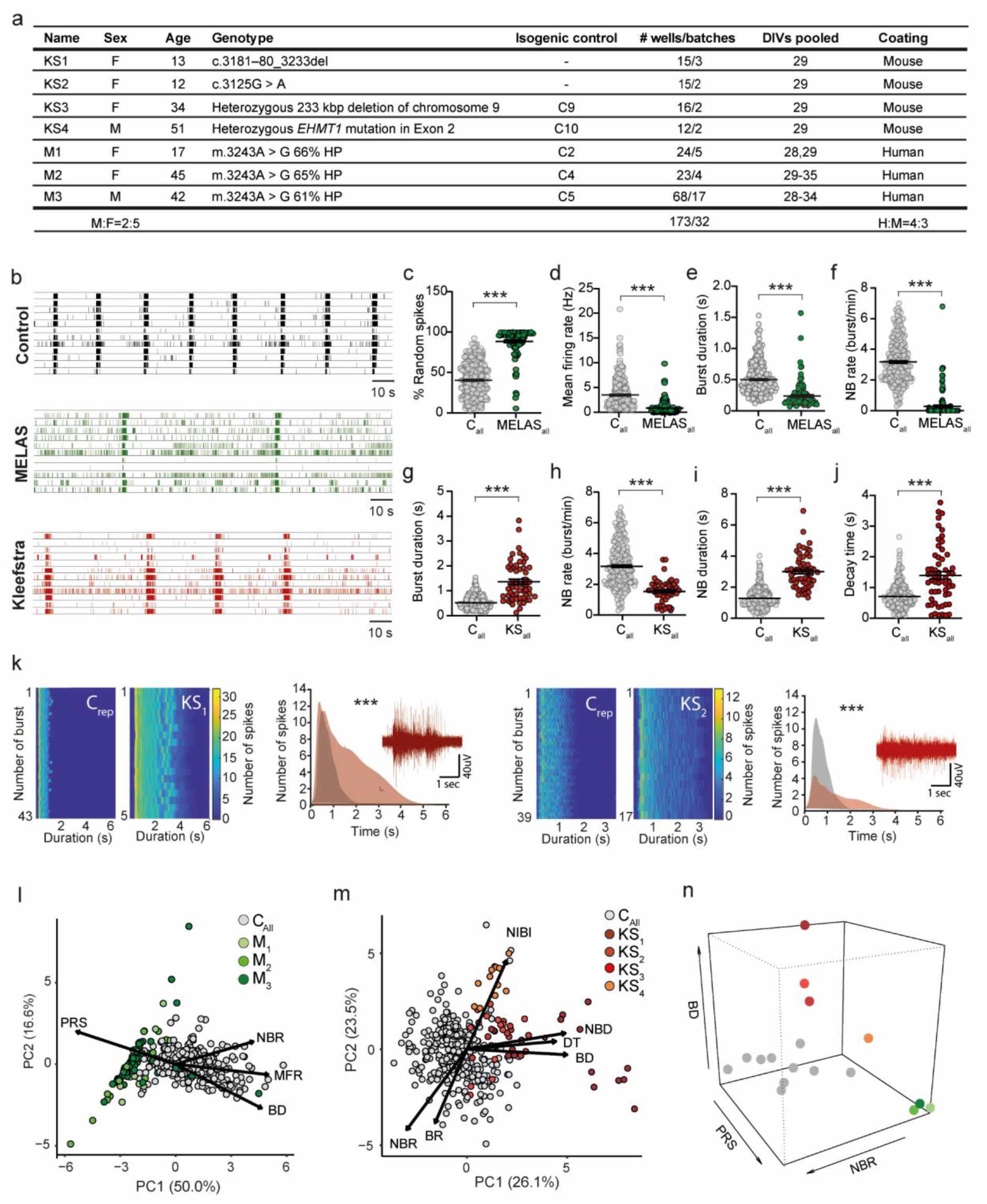
MEAs pose a reliable platform for genotype-phenotype correlations. (**a**) Information regarding the 7 patient lines included in this study. Isogenic controls represent the lines made from the same founder somatic cell line. Number of wells represent total number of wells recorded for that line between DIV27-35, including the number of batches. Some batches overlap between lines. (**b**) Representative raster plots showing 3 minutes of electrophysiological activity from control, MELAS and KS patient lines. (**c-f**) Graphs showing the values of four MEA parameters including (**c**) PRS, (**d**) MFR, (**e**) BD, and (**f**) NBR for control and MELAS neuronal networks. (**g-j**) Graphs showing four MEA including (**g**) BD, (**h**) NBR, (**i**) NBD and (**j**) DT for control and KS neuronal networks. Mann Whitney U test with Bonferroni correction for multiple testing was used to compare between patient lines and their isogenic controls (**Table S6**). (**k**) Representative network burst alignment from one recording of a representative control and KS1, and a representative control and KS2. Inset: extracted burst shape and representative raw trace of a network burst (Sample size n for C representative C_6_=58, C_9_=12, KS_1_=15, KS_2_=15, multiple *t*-test on bins using Holm-Sidak method, p <0,0001 for both comparisons). (**l**) PCA plot on 7 MEA parameters, showing parameters that explain the differences in network behavior between C_all_ (278 wells from 10 control lines) and M1-3. (**m**) PCA plot on 12 MEA parameters, showing parameters that explain the differences in network behavior between C_all_ and KS1-4. (**n**) 3D scatter plot showing PRS, BD, and NBR for all MELAS (green), KS (red), and control lines (grey). *p* = 0.01 ** and *p* = 0.001 ***. DIV = days in vitro, MFR = mean firing rate, PRS = percentage of random spikes, BD = mean burst duration, NBR = network burst rate, NBD = network burst duration, DT = decay time. All means, *p*-values and used statistic tests are reported in supplementary Table S6.

Neuronal networks from MELAS patients showed a different network phenotype compared to the control pool (**Fig. 3b**). In line with previous findings (Hsieh et al. 2005, Gunnewiek et al. 2020), the phenotype was mainly explained by a strong reduction in level of spiking and network bursting activity, together with an increased percentage of random spikes (**Fig. 3c-f**). In addition to previously published data, we found that neuronal networks derived from MELAS patients exhibited burst with a shorter duration compared to the control pool (**Fig. 3e**). We did not observe any difference in burst shape, rise time (RT) and decay time of MELAS patient network bursts compared to controls (**Fig. S4a**). Neuronal networks derived from MELAS patients did show a lower level of correlation (C_peak_) and synchronization (C_0_) among all channels compared to controls (**Fig. S4b, Table S1**). Furthermore, despite a comparable number of functional connections amongst electrodes in control and MELAS patient neuronal networks, we observed that the connections between MELAS neurons were weaker (**Fig. S4c**). PCA confirmed MELAS patient networks clustered separately from controls (**Fig. 3l**).

Next, we compared neuronal networks from patients with KS to our total control pool and uncovered a significantly different network phenotype (**Fig. 3b**). In line with previously published findings (Frega, Linda et al. 2019), the KS phenotype was mainly characterized by a lower frequency of (network) bursts with a longer duration (**Fig. 3g-i**). In addition, we found a different network burst shape and an increased decay time in KS neuronal networks (**Fig. 3j,k**). Furthermore, KS neuronal networks showed lower level of synchronicity and correlation and weaker connections between neurons as compared to controls (**Fig. S4d,e**). We observed that differences from controls were more pronounced in KS_4_ as compared to the other KS lines (**Fig. S4f**). PCA confirmed that KS neuronal networks clustered away from controls based on these parameters (**Fig. 3m**).

In conclusion, the neuronal network phenotypes of MELAS and KS lines differed from controls on dinstinct parameters. Indeed, MELAS and KS samples cluster away from controls, but also clearly cluster away from each other (**Fig. 3n**). This ability to distinguish two NDDs based on their neuronal network phenotypes demonstrates that the MEAs system is a adequate platform for disease-specific phenotyping.

### Comparing patients with isogenic controls reveals a more detailed phenotype

Isogenic hiPSC lines are increasingly used to improve identification of genotype-phenotype correlations. We compared data from three MELAS mosaic patient-control isogenic sets, one KS mosaic patient-control set and one KS CRISPR-Cas9 engineerd isogenic set. The difference between each MELAS and KS isogenic set was explained by the same parameters as when patient lines were compared to all control lines (**Fig. 3, Fig. 4**). We additionally found that the difference between patient and control lines was larger for isogenic sets compared to all lines combined, as indicated by the higher variance explained by disease status (**Fig. 3l, m** and **Fig. 4g, n**).

**Figure 4.**
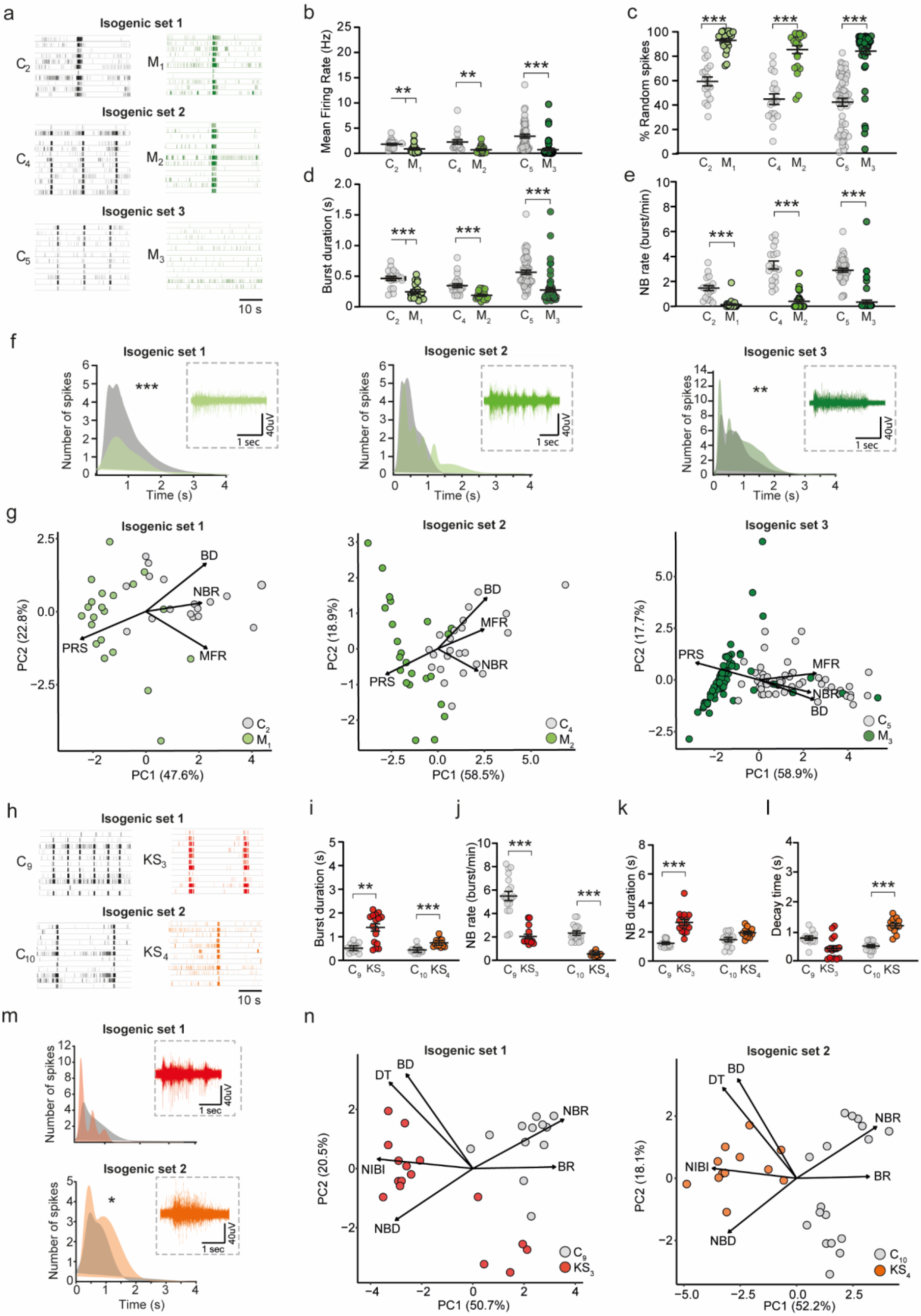
Characterization of isogenic control and patient networks. (**a**) Representative raster plots showing 1 minute of activity from 3 control (C_2_, C_4_, C_5_)-MELAS (M_1-3_) isogenic sets. (**b-e**) Comparison of the MEA parameters (**b**) MFR, (**c**) PRS, (**d**) BD, and (**e**) NBR for each corresponding MELAS isogenic patient-control set. (**f**) Burst shape and representative raw trace of a network burst from C_2_ and M_1_, C_4_ and M_2_ and C_5_ and M_3_ (Sample size n for C_2_ = 15, C_4_ = 23, C_5_ = 55, M_1_ = 22, M_2_ = 8 and M_3_ = 7, multiple *t*-test on bins using Holm-Sidak method, C_2_ vs M_1_ *p* <0.001, C_5_ vs M_3_ *p* = 0.00021) (**Table S7**). (**g**) PCA plots on 7 MEA parameters for MELAS isogenic patient-control sets C2 and M1, C4 and M2 and C5 and M3 showing MEA parameters affected in MELAS. (**h**) Representative raster plots showing 1 minute of activity from 2 control (C_9-10_)-KS patient (KS_3-4_) isogenic sets. (**i-l**) Comparison of the MEA parameters (**i**) BD, (**j**) NBR, (**k**) NBD, and (**l**) DT for each corresponding KS isogenic patient-control set. (**m**) Burst shape and representative raw trace of a network burst from C_9_ and KS_1_, C_10_ and KS_4_ (sample size n for C_9_ = 12, C_10_ = 17, KS_3_ = 16, KS_4_ = 12, multiple *t*-test on bins using Holm-Sidak method) (**Table S7**). (**n**) PCA plot on 12 MEA parameters for KS isogenic patient-control sets C_9_ and KS_3_ and C_10_ and KS_4_ showing MEA parameters affected in KS. *p* = 0.05*, *p* = 0.01 ** and *p* = 0.001 ***. DIV = days in vitro, MFR = mean firing rate, PRS = percentage of random spikes, BR = mean burst rate, BD = mean burst duration, BSR = Burst spike rate, IBI = inter-burst interval, NBR = network burst rate, NBD = network burst duration, NIBI = Network burst IBI, CV_NIBI_ = coefficient of variation of all NIBI’s representing the regularity of the NB, RT = Rise time, DT = decay time. All means, *p*-values and used statistic tests are reported in supplementary Table S7.

For some parameters, we observed smaller differences in one MELAS set (i.e. isogenic set 1) as compared to the other isogenic sets (**Fig. 4a-e**). This was mainly driven by a difference in the isogenic control (C_2_) compared to all controls, rather than a less pronounced MELAS phenotype (**Fig. S1f-o**). While comparing isogenic sets, we found a significant difference in the network burst shape of two MELAS isogenic patient-control sets (**Fig. 4f**), a phenotype that was not distinguished when comparing MELAS patients to control pool. Some line-specific differences were also found in the MEA parameters explaining the KS network phenotype. Whereas the decay time was not affected in KS_3_ compared to its isogenic control, and only a trend was observed in network burst duration for KS_4_, these parameters were shown to be affected in all other KS lines (**Fig. 4k-m, Fig. S4g**). Post-hoc power calculation revealed that the parameters that explained the MELAS and KS phenotypes reached a power higher than 0.95 (**Table S9**), demonstrating the validity of our results. When performing a priori power calculation on each patient-control isogenic set or all controls compared to all patient lines, we found that a minimum of 12 wells per line should be included in the analysis to observe a patient phenotype on multiple MEA parameters (**Table S9**).

To conclude, disease phenotypes are generally consistent between different lines from patients with the same disorder, even though some line-specific differences can be observed. This was also true when comparing MELAS and KS lines to their corresponding isogenic controls, highlighting the importance of using multiple patient lines to uncover the full phenotypic spectrum. Nevertheless, we show that isogenic patient-control sets uncover more pronounced differences, emphasizing the advantage of using isogenic sets to uncover patient phenotypes.

## Discussion

MEAs have been used for more than four decades to study neuronal network dynamics *in vitro* (Gross et al. 1982). The recent advances in hiPSC technology enabled the study of neuronal network functionality in healthy and diseased human neurons. Despite the increasing popularity of MEAs for disease phenotyping, there is little insight into the range of variability between human control neuronal networks and the conditions that influence this variability. In this study, we performed a meta-analysis of, to our knowledge, the largest dataset of hiPSC-derived *Ngn2*-induced excitatory neuronal networks on MEAs to describe a standard for control neuronal network signatures.

We uncovered that data recorded from networks derived from ten different healthy subjects clustered together in PCA, regardless of whether they were cultured by different researchers over the course of years, and independent of the sex and age at fibroblast biopsy. These control neuronal networks were very comparable because our lab adhered to a strict set of guidelines (**Fig. 2k**). First, networks could only be pooled in the time window between DIV 27-35 because neuronal networks generated by *Ngn2*-overexpression in our lab presented stable network activity at this stage. However, many factors can influence the timing of this stable network activity that are often controlled by the user, which can induce a shift of this stable period. For example, this stable period of network activity can significantly differ between neuronal differentiation protocols, as *Ngn2*-neurons mature significantly faster than neurons generated using small molecule supplementation protocols (Lomvardas et al. 2016, Mertens et al. 2016). In addition, the developmental trajectory in neuronal networks grown on different coating reagents can differ. While neuronal networks grown on human and mouse laminin showed no difference after DIV 28, cultures grown on human laminin displayed slower maturation (Flemming 1882, Hyysalo et al. 2017). When one wants to pool and compare data, one must first define the stable developmental period depending on each protocol, before pooling and comparing data.

Second, as hiPSC culture practices and differentiation protocols consist of many steps, small differences in handling cells between batches or by different researchers can accumulate over time into significantly different outcomes, and thus introducing variation in the data (Luger et al. 1997, Volpato et al. 2020). For example, we showed that astrocyte isolation batch and MEA batch influenced the variability of the data. Moreover, our power calculation revealed that a minimum of 12 wells should be analyzed. We advise to divide these wells across multiple batches (i.e. at least two) to account for variation in MEA batch. We showed that neuronal networks with low cell densities or an altered distribution of cells across the electrodes display activity that was not comparable to neuronal networks that were cultured at optimal conditions. It is advised to critically look at the cell density and distribution on the electrode grid in conjunction with the corresponding data, and exclude wells with low density to obtain robust results.

Third, to extract biological phenotypic signatures present in the neuronal networks on MEAs it is essential to perform an accurate data analysis and include multiple parameters describing the neuronal activity. The choice of the analysis settings for extraction of data on MEA parameters can largely influence the results. Indeed, we showed that neuronal networks in which not all network burst were detected correctly significantly differed from the control pool. To accurately detect different phenotypic signatures, it is possible that the analysis settings should be fine-tuned or changed. For example, network burst exhibited by patient-derived neuronal networks might be wrongly detected with commonly used settings, since these were conventionally chosen based on how network bursts appeared in control networks, mainly from rodent origin. Moreover, it is possible that the observed phenotype cannot be described using any of the commonly used parameters, and new parameters should be introduced to capture these signatures. Indeed, we showed that the extraction of additional parameters from MEA data revealed previously unseen phenotypes. The burst shape and correlation analysis uncovered previously unreported differences in both KS and MELAS neuronal networks. Attention should be payed when determining which parameters will be used to describe the network phenotype. We have also determined the variability of the MEA parameters since stable parameters are the most trustworthy to identify a disease phenotype. In the literature differences in neuronal network phenotypes are sometimes solely based on mean firing rate (MFR) (Heitz 1928, Luger et al. 1997, Wainger et al. 2014, Sadakierska-Chudy et al. 2015, Chailangkarn et al. 2016, Lu et al. 2016), as it is easily extracted from the data. Interestingly, our data shows that the MFR is one of the most variable parameters. In addition, it only describes the general level of neuronal network activity and it is furthermore largely dependent on cell density. The MFR should therefore be interpreted with caution when solely used to describe a phenotype. Not only the MFR, but also the inter burst interval (IBI), CV_NIBI_, RT, C_0_ and link weight were variable parameters. Similar to the MFR, the IBI is dependent on cell density, as the probability to detect a burst (i.e. 4 spikes in close proximity) is becoming smaller with lower densities. A similar principle holds true for the RT, as a low cell density can affect the number of spikes composing a network burst, and thereby affects the network burst shape. Regarding the connectivity measures, the observed variability might be influenced by low number of recording electrodes and thus more stable results can be obtained using high-density MEAs. For this reason, MEA parameters with high variability can be used to determine a patient phenotype, but multiple supporting MEA parameters should be included to describe the neuronal network characteristics.

It is generally accepted that mutations in genes causing NDDs all result in some form of neuronal circuit dysfunction. When we compared our control and disease-specific neuronal networks, we found a strong segregation between control and MELAS or KS neuronal networks, as well as between MELAS and KS networks. We found that KS and MELAS neuronal networks were distinguished by different MEA parameters. Literature has shown that even when investigating different genes associated with the same NDD, neuronal networks showed a similar phenotype, albeit characterized by an individual set of parameters depending on investigated gene. For example, we previously identified that rat cortical networks deficient for the KS spectrum genes *Ehmt1, Mll3, Mbd5* and *Smarcb1* all displayed hyperactive neuronal networks. However, whereas EHMT1- and SMARCB1-deficient networks showed a significantly higher MFR, MLL3-deficient networks showed a higher NBR (Davis et al. 2018, Frega et al. 2020). Likewise, a recent study that investigated *Ngn2*-induced neuronal network behavior of 12 autism spectrum disorder (ASD) patients revealed hyperactive neuronal networks specifically from a patient with *CNTN5* and an CRISPR-Cas9 engineered line with an *EHMT2* mutation. In neuronal networks with a *CNTN5* mutation, the network phenotype exhibited differences in NBR, whereas in networks with an *EHMT2* mutation this was the case for the MFR (Deneault et al. 2019). Together, this strengthens the evidence that early disease-associated network phenotypes can be revealed using MEAs, and that hiPSC-derived neurons are a powerful model to study genotype-phenotype correlations.

Although control neuronal networks were robust enough to perform disease phenotyping, it must be noted that we still observed significant variation between different control lines. This variation likely reflects normal variation in the general population (Bjornsson 2015, Germain et al. 2017). Gene expression and DNA methylation in hiPSC lines varies significantly amongst healthy donors, of which common genetic variation is the main driver (Reinke et al. 2003, Zhao et al. 2005, DeBoever et al. 2017, Germain et al. 2017, Kilpinen et al. 2017, Mossink 2020). Indeed, previous literature uncovered that the heterogeneity within 25 different hiPSC lines on a transcriptional level was due to differences in genetic background (Rouhani et al. 2014). Adding to this, variation between cell lines from different individuals can be more pronounced in differentiated cell types (Hassan et al. 2006, Banovich et al. 2018), and differentiation efficiency of hiPSCs into neurons itself can also contribute to variation seen between lines (Hu et al. 2010). We speculate that this difference in common genetic variation and differentiation efficiency can result in small variations on a functional level, similar to what was observed here.

In line with minor variations that we see in control neuronal networks, we observed a similar variability in patient-derived lines. We detected patient line-specific differences, especially for KS neuronal networks that all carry heterozygous mutations in *EHMT1.* Differences in network burst characteristics generally define the phenotype of these KS lines, but some lines showed more pronounced differences than others. While we cannot rule out a patient specific component, our functional analysis revealed that at least sex or age did not interfere with identification of phenotype on MEA. Indeed, the disease effect was larger than the age or sex effect, since KS and MELAS neuronal networks clustered away from controls independently from the donor background. However, to correct for line specific differences and variability, multiple lines from different individuals should be used.

Another strategy to reduce the variability between patient and control neuronal networks is the use of isogenic patient-control sets, since these are not influenced by differences in genetic background (Bowman et al. 2015, Bassett 2017, Germain et al. 2017). We observed a more pronounced phenotype when isogenic control and patient lines were used. There are two routes to generate isogenic patient-control sets, which can be used complementary to each other (Ruthenburg et al. 2007, Bassett 2017). One can either derive a control line from a patient by correcting the patient mutation, or introduce a patient mutation in a control line (Wang et al. 2011, Bassett 2017). It is also possible to derive both a healthy and diseased hiPSC line from a mosaic carrying patient, which we here show for both C_9-10_ and KS_3-4_ as well as C_2_, C_4_, C_6_ and M_1-3_ hiPSC lines. While using isogenic patient-control sets derived from patient background provides evidence that the particular gene contributes to the patient phenotype, it remains unknown if this gene alone is sufficient to cause disease or whether there is any unidentified genetic variation in the patient background that contributes to the phenotype. In our dataset, one MELAS isogenic patient-control set showed a less pronounced neuronal network phenotype. This was caused by a slightly altered network phenotype of control line C_2_ compared to the other controls, whereas the network phenotype of this specific MELAS line was similar to the other MELAS lines. It could be that the individual’s genetic background influences the control phenotype, despite the absence of the m.3243A>G mutation. Therefore, it is worth to also investigate if introducing a patient mutation in a healthy control line produces a similar phenotype to show a more direct genotype-phenotype relationship.

In summary, we here provide a set of guidelines to reduce the variability in neuronal network recordings on MEA, which are summarized in **Fig. 2k**. We project that if cultures are handled according to these guidelines, our control dataset can be used as a reference database to determine the performance of *Ngn2*-induced control lines. An extensive list of literature has shown that network parameters can differ between different sources, neuronal differentiation protocols, or species (Heikkilä et al. 2009, Napoli et al. 2016, Odawara et al. 2016, Hyysalo et al. 2017). While we expect that other neuronal model systems will show network parameters in a different range than reported here, the guidelines that we propose can nevertheless be applied to other neuronal model systems as well. First, we recommend that when a MEA experiment is designed, at least 12 independent wells divided over at least two neuronal preparations should be used. Furthermore, as we identified a clear MEA- and astrocyte-batch effect on neuronal phenotype, it is advised to compare the different experimental conditions using astrocytes from the same batch, and evenly distribute conditions among different MEA batches. We advise to use stringent exclusion criteria, including exclusion of wells with poor health or without an uniform distribution of cells based on visual inspection. Furthermore, control neuronal networks showing very low levels of activity (i.e. MFR <0.1 spike/s, MBR < 0.4 burst/min, active channels < 80 % and NBR < 1 burst/min) should be excluded. Moreover, we recommend extracting multiple parameters from the recording to describe the neuronal network phenotype, and highly discourage the use of only one MEA parameter to describe a patient phenotype. To correct for cell line specific variation in control hiPSC lines, and to uncover the full phenotypic spectrum of patient lines, at least three patient and control lines should be used to describe the neuronal network phenotype. Additionally, to overcome general variation in the control and patient cohort, isogenic patient-control sets can be used to further characterize the phenotype in depth. Following these guidelines, MEAs are a valuable tool to describe the neuronal network phenotypes in hiPSC-derived neuronal networks.

## Methods

### Control hiPSC line origin and generation

All hiPSC lines in the studies used to generate this dataset were obtained by reprogrammed skin fibroblasts. We have used ten hiPSC control lines in total, of which three are commercial control lines (C_1_, C_6_, C_10_) and two are control lines generated in house (C_3_ and C_8_). Specifically, control line C_1_ was derived from a curated control cell line (Jones et al. 1998, Bestor 2000, Okita et al. 2011, Kondo et al. 2017). This control line originated from a 36-year old female, reprogrammed using episomal vector-based reprogramming of the Yamanaka transcription factors *Oct4, c-Myc, Sox2* and *Klf4* (Nan et al. 1998, Takahashi et al. 2006), showing no karyotypical malformations. Control line **C_6_** was derived from a 30-year old male (Williams et al. 2010, Talkowski et al. 2011, Miyaoka et al. 2014, Mandegar et al. 2016) and reprogrammed using episomal vector-based reprogramming of the Yamanaka factors and was tested for genetic integrity using SNP assay (Bird 2008, Frega et al. 2019). Control line **C_7_** was derived from a 41-year-old-female and reprogrammed using the Simplicon™ reprogramming kit (Millipore). Overexpression of the Yamanaka factors was introduced by a non-integrative, non-viral one-step transfection. Genomic stability was checked using STR analysis. Control line **C_8_** was derived from a 9-year old male and reprogrammed using retroviral-vector based reprogramming of the Yamanaka factors and tested for pluripotency and genomic integrity based on SNP arrays.

To illustrate that the model that we use is stable enough to uncover patient specific phenotypes, we included both KS patient and MELAS patient lines, as well as isogenic patient-control sets. The remaining five hiPSC control lines corresponded to isogenic patient-control sets, as shown in **Fig. 1a and Fig. 3a** (C_2-5_, C_7_ and C_10_). The KS isogenic patient-control hiPSC set consisting of control line **C_9_ and KS_3_** was derived from a 34-year old female with a mosaic heterozygous 233 kbp deletion of chromosome 9, including the *EHMT1* and *CACNA1B* gene, diagnosed with KS (Tyagi et al. 2016, Frega et al. 2019). Both clones were reprogrammed using retro-viral vectors expressing the Yamanaka factors and tested for genomic integrity based on SNP array (Krogan et al. 2003, Frega et al. 2019). The second KS isogenic patient-control hiPSC set, consisting of control line **C_10_ and KS_4_** was previously derived from a healthy 51-year old male and reprogrammed using expression of Yamanaka factors by non-integrating Sendai virus. KS_4_ was generated using CRISPR/Cas 9 technology to induce a heterozygous *EHMT1* mutation to mimic KS, as described previously, producing an isogenic set (Tyagi et al. 2016, Frega et al. 2019). Both lines were tested for genomic integrity based on SNP array (Tyagi et al. 2016, Frega et al. 2019). In addition, we included two KS patient hiPSC lines, **KS_1_ and KS_2_**, which were previously characterized and derived from a 13-year old and a 12-year-old female, respectively, diagnosed with KS (Cremer et al. 2001, Frega et al. 2019). HiPSC clones were obtained by reprogramming of retroviral expression of the Yamanaka factors.

The MELAS syndrome isogenic patient-control hiPSC sets consisting of control line **C_2_, C_3_ and C_4_ and patient lines M_1_ and M_2_** were generous gifts form Esther-Perales Clemente and Timothy Nelson. C_2_ and M_1_ were derived from a 17-year-old female, and C_4_ and M_2_ form a 45-year-old female. Both donors were diagnosed with MELAS syndrome, harboring the pathogenic variant m.3243A>G. Reprogramming of fibroblasts to hiPSCs resulted in clones with varying levels of m.3243A>G heteroplasmy (Lieberman-Aiden et al. 2009, Dixon et al. 2012, Perales-Clemente et al. 2016, Gunnewiek et al. 2020). Control lines C2 and C_4_ originated from hiPSC clones with confirmed 0% heteroplasmy, while M_1_ and M_2_ had 66% of m.3243A>G heteroplasmy and 80% heteroplasmy, respectively. Control line C_3_ was derived from a 30-year-old male with MELAS syndrome and was confirmed to have 0% heteroplasmy upon reprogramming to hiPSCs. All lines were reprogrammed using CytoTune-iPS Sendai Reprogramming Kits according to manufacturer’s instructions (Invitrogen, A13780-02, A16517, A16518) and were previously characterized and had a normal karyotype (de Laat et al. 2013). MELAS syndrome isogenic patient-control hiPSC set, consisting of control line **C_5_ and M_3_** was derived from a 42-year-old male with MELAS syndrome with a pathogenic variant m.3243A>G. Reprogramming through lentiviral transduction of the Yamanaka transcription factors resulted in hiPSC clones with varying levels of m.3243A>G heteroplasmy (Phillips-Cremins et al. 2013, Gunnewiek et al. 2020). C5 originated from an hiPSC clone with confirmed 0% of m.3243A>G heteroplasmy and M3 originated from an hiPSC clone with 70% heteroplasmy. Both C_5_ and M_3_ were previously characterized and had normal karyotypes (Rao et al. 2014, Gunnewiek et al. 2020).

All generated hiPSC clones were tested for pluripotency markers (OCT3/4, SOX2 and NANOG) using immunocytochemistry and q-PCR. HiPSCs were cultured on E8 Flex basal medium (ThermoFisher scientific #A2858501) supplemented with primocin (0.1 μg/ml, Invivogen #ant-pm-1), puromycin (0.5 μg/ml) (Sigma-Aldrich #P9620) and G418 (50 μg/ml) (Sigma-Aldrich #A1720) at 37 °C/5% CO2, on either human recombinant laminin LN521 (Biolamina #LN521-02) or Matrigel (Corning #356237) coated plates. Medium was refreshed every two days and cells were passaged approximately every three days using ReLeSR (ReLeSR, Stem Cell Technologies #05873), an enzyme free passaging reagent.

### Neuronal differentiation and culture

HiPSCs were differentiated into upper layer excitatory cortical neurons by doxycycline inducible expression of the neuronal transcription factor Neurogenin 2 (*Ngn2*) (Xiao et al. 2011, Zhang et al. 2013), according to a previously published protocol (Vietri Rudan et al. 2015, Frega et al. 2017). To generate single cells, *rtTA/Ngn2* positive hiPSCs were detached by incubating accutase (Sigma-Aldrich #A6964) at 37°C/5%CO_2_ and resuspended in E8 basal medium (ThermoFisher #A15170-01), supplemented with primocin (0.1 μg/ml), RevitaCell (ThermoFisher #A2644501) (10 μg/mL) and doxycycline (Sigma-Aldrich #D9891) (4 μg/mL) to induce TetO gene expression. Cells were plated at a density of 20.000 cells per MEA well (600 neurons/mm^2^), which were pre-coated with poly-L-ornithine hydrobromide ((PLO), Sigma-Aldrich #P3655-10MG) (50 mg/mL) and, depending on experiment, either human recombinant laminin LN521 (5 mg/mL) or laminin from Engelbreth-Holm-Swarm murine sarcoma basement membrane ((mouse laminin), Sigma-Aldrich #L2020) (20 μg/mL). At DIV1 medium was changed using filtered DMEM/F12 supplemented with primocin (0.1 μg/ml), doxycycline (4 μg/mL), 1% N-2 supplement (ThermoFisher #17502-048), 1% MEM non-essential amino acid solution (Sigma-Aldrich #M7145), Neurotrophin-3 ((NT3), Promokine, #C-66425) (10 ng/mL), recombinant human brain derived neurotrophic factor ((BDNF), Promokine #C-66212) (10 ng/mL) and mouse laminin (0.2 μg/mL). At DIV2, rat embryonic astrocytes were added in a 1:1 ratio to support neuronal maturation and viability (Schwalie et al. 2013, Frega et al. 2017). The medium was changed at DIV3 to filtered Neurobasal medium (ThermoFisher #21103-049) supplemented with primocin (0.1 μg/ml), B-27 (ThermoFisher #17504044) (20 μg/mL), GlutaMAX (ThermoFisher #35050061) (10 μg/mL), doxycycline (4 μg/mL), NT3 (10 ng/mL), BDNF (10 ng/mL), and Cytosine β-D-arabinofuranoside ((Ara-C), Sigma-Aldrich #C1768) (2 μM), to remove proliferating cells from the culture. From DIV 5-DIV 9 ~50% of the Neurobasal medium supplemented with B-27, GlutaMAX, Pen/Strep, doxycycline, NT3 and BDNF, was refreshed every 2 days. From DIV 9-21 onwards the neurobasal medium was in addition supplemented with 2.5% fetal bovine serum (FBS, (Sigma-Aldrich #F7524)) to support astrocyte viability. All neuronal cultures were kept in incubation at 37°C/5%CO_2_. Control lines C1, C6 and C7 were partly cultured in the absence of doxycycline from DIV13 onwards. No significant effect between wells cultured with and without doxycycline were found, therefore all data for these respective lines is pooled (data not shown).

### MEA recordings and data analysis

To record spontaneous network activity, multiwell-MEAs were used that consisted of 24 individual wells (Multichannel systems, MCS GmbH, Reutlingen, Germany). Each well was embedded with 12 electrodes with a diameter of 30 μm, spaced 300 μm apart. The activity of neuronal networks growing on MEAs was recorded for 10 minutes (after an acclimatization period of ten minutes) in a recording chamber that was maintained constant at 37°C/95% O_2_/5% CO_2_. The raw signal was sampled at 10 kHz and filtered with a high-pass 2^nd^ order Butterworth filter with a 100 Hz cut-off frequency and a low-pass 4^th^ order Butterworth filter with a 3500 Hz cut-off frequency. The noise threshold for individual spike detection was set at ±4.5 standard deviations.

#### Data analysis using Multiwell-Analyzer

Off-line data analysis was performed using Multiwell-Analyzer software (Multichannel systems) that permitted the extraction of spike-trains, and either a custom-made in-house code developed in MATLAB (The Mathworks, Natick, MA, USA) or a software package called SPYCODE (Bologna et al. 2010, Cournac et al. 2015, Mourad et al. 2016, Frega et al. 2019), which both allowed the extraction of parameters describing the spontaneous network activity.

The mean firing rate (MFR) (Hz) was calculated for each well individually by averaging the firing rate of each separate channel by all the active channels of the well. Bursts were detected using the Multiwell analyzer build-in burst detection algorithm. The algorithm was set to define bursts if 4 spikes were in close proximity with a maximum of 50 ms inter spike interval (ISI) to start a burst, and a maximum of 50 ms ISI to end a burst, with a minimum of 100 ms inter burst interval (IBI).

Network bursts were defined when at least 50% of all channels simultaneously displayed a burst. In rare cases, we observed synchronous events which were not properly identified as network bursts (**Fig. 2g**). It was however clear that the observed pattern of the synchronous events was similar to control networks in which network bursts were properly detected. When the network burst detection was insufficient the network bursts detection was decreased down to, but not further than at least 25% of all channels participating in the network burst.

The percentage of random spikes (PRS) was defined by calculating the percentage of spikes that neither belonged to a burst nor a network burst. The IBI, and network burst IBI (NIBI) were calculated by the subtraction of the time stamp of the beginning of each burst or network burst, respectively, from the time stamp of the ending of the previous. To calculate the IBI, the mean IBI per channel was calculated and averaged across all channels. The NIBI is calculated by averaging the found NIBIs in one well. To explain the regularity at which a network burst occurs, the coefficient of variation (CV) was calculated by calculating the standard deviation of all NIBIs and dividing it by the mean of all NIBIs.

#### Analysis of the average burst shape

The generation of average burst shapes for each control was performed using adapting scripts and functions implemented in MATLAB (Melé et al. 2016, Van De Vijver et al. 2019). Spike trains containing all events detected in the 12 electrodes were binned. Network burst were detected from a spike train containing the events present in all channels by using an inter-spike interval threshold (30 ms). All network bursts that have a duration of less than 100 ms were removed from the detection. For each well, the average burst shape was calculated by aligning all the network bursts detected to the longest network burst and by summing all spikes (bin: 10 ms). Then, we calculated the average burst shape for each line by averaging all the histograms of the individual wells. We fitted the curves with Gaussian distribution to obtain smoother profiles using the build in fit function. The rise and decay times were calculated as the absolute difference between the time at 20% and 80% before and after the peak, respectively. Slopes of rise and decay time were subsequently calculated using the build in polyfit and polyval functions.

#### Correlation analysis

We obtained connectivity matrix for each culture by applying the Filtered Cross-Correlation algorithm (Gorkin et al. 2014, Pastore et al. 2018) available in the free software SPICODYN (Pastore et al. 2018, Bastle et al. 2019). Cross correlograms were obtained using a bin size of 0.1 ms and a correlation window of 150 ms. Matrices were then analyzed using custom script in MATLAB. We evaluated the C_peak_ (i.e. maximum value of the-cross-correlation function), C_0_ (i.e. value of the cross-correlation function in the central bin), total number of links and the average weight of those links for each culture.

To guarantee sufficient experimental replicates, we included experiments with a minimum of 12 wells per hiPSC line measured across at least two independent batches. Control neuronal networks showing a MFR <0.1 Hz and burst rate <0.4 bursts per minute were excluded from analysis. Wells were excluded from analysis if they did not have network bursts at DIV 27. Furthermore, wells that displayed insufficient quality, for example a low density of cells or cell clumping, were discarded. All experiments, excluding experiments where we investigated neuronal network development over time, were carried out during a one-week time interval, spanning DIV 27 till DIV 35. Since our results, and previous research has shown that network burst parameters are stable from DIV 27 onwards, data from DIV 27 till DIV 35 were pooled (Bjornsson 2015, Frega et al. 2019). When analyzing multiple developmental time points of one MEA batch, we determined the network burst detection settings at the latest DIV and kept these settings throughout the analysis, working our way backwards to the earliest DIV. Wells in which a reduction of network parameters was observed were also excluded.

### Statistical analysis

Data were analyzed using Prism GraphPad 8 (GraphPad Software, Inc., CA, USA). We ensured normal distribution using a Kolmogorov-Smirnov normality test. To determine statistical significance *p*-values <0.05 were considered to be significant. Statistical analysis on all control lines in **Fig. S1f-o, Fig. S2b, c, Fig. S3b-e and Fig. S4f-g** (for KS1 and KS2) were performed using a Kruskal-Wallis ANOVA with post hoc Dunn’s correction for multiple testing or One-way Anova with Tukey correction or multiple testing depending on the distribution of the data. When comparing means of two variables at one individual timepoint we analyzed significance between groups by means a Mann-Whitney-U-Test (**Fig. 3c-j, Fig. 4b-e and i-l, Fig. S4f** (for KS3 and KS4), and if applicable, corrected post hoc for multiple testing using the Bonferroni method. Statistics on histograms was performed using Multiple *t*-test on bins using Holm-Sidak method (**Fig. 3k and Fig. 4f, m**). Data are presented as mean ± standard error of the mean (SEM) if not differently specified. Means and *p*-values are reported in **Table S3-10**. To check the variability in the dataset we calculated the coefficient of variation (CV) on each parameter independently for all control lines (**Fig 1j, Fig S3g and Table S2**).

### Principal component analysis

Principal component analysis (PCA) was performed on data from 12 MEA parameters (MFR, PRS, BR, BD, BSR, IBI, NBR, NBD, NIBI, CV_NIBI_, RT, and DT, see table 1) for all control samples (n_wells_ = 278, N_plates_ = 47). PCA was performed using the prcomp function from stats R package (v3.6.1.) on standardized (z-score scaled) data. Separately, PCA was performed for controls including samples from either KS (n_wells_ = 58, N_plates_ = 9) or MELAS (n_wells_ = 112, N_plates_ = 23) neurons. PCA on KS versus control samples was performed on data from all 12 MEA parameters. PCA was performed for each KS isogenic patient-control set separately as well, using the same MEA parameters. PCA on MELAS versus control samples was performed on data from 7 MEA parameters (MFR, PRS, BR, BD, BSR, IBI, and NBR), since many samples did not show network bursts (i.e. NBD, NIBI, CV_NIBI_, RT, and DT could not be calculated). PCA was performed for each MELAS isogenic patient-control set separately as well, using the same MEA parameters.

PCA’s in **Fig. 2i, j** were performed for samples from line C1 separately to test the effect of astrocyte batch and MEA plate, using data from all 12 MEA parameters. Samples from C1 were measured using three different astrocyte batches. Within three astrocyte batches, samples were measured on 2-3 different MEA plates. PCA was performed on all C1 samples (n_wells_ = 27, N_plates_ = 7) to check the effect of astrocyte batch, as well as on samples from these three individual astrocyte batches separately (n_wells_ = 10, N_plates_ = 3; n_wells_ = 9, N_plates_ = 2; n_wells_ = 8, N_plates_ = 2) to check the effect of MEA plate batch.

Furthermore, PCA in **Fig. S1p** was performed for samples from line C6 that were measured up to DIV62, to check whether samples from early DIVs cluster away from samples at a higher DIVs. This was performed for two MEA plates separately, for which data was available between DIV14-56 (n_wells_ = 7) and DIV24-62 (n_wells_ = 4), using data from 7 MEA parameters (MFR, PRS, BR, BD, BSR, IBI, and NBR), since at an early time point (almost) no network bursts are detected (i.e. NBD, NIBI, CV_NIBI_, RT, and DT could not be calculated).

Results from PCA are depicted in figures showing data points plotted on PC1 vs PC2 including the variance explained per principal component, for all different analyses performed. Samples are labeled according to variable of interest per figure. PCA figures were generated using the ggplot function from ggplot2 R package (version 3.2.1). A 3D scatter plot was made for all control (n_wells_=278, N_plates_ = 47), Kleefstra (n_wells_ = 58, N_plates_ = 9) and MELAS (n_wells_ = 112, N_plates_ = 23) samples together showing PRS, BD, and NBR using the scatter3d function from scatterplot3d R package (version 0.3-41).

### Animals

The rodent astrocytes used in this study were derived from embryonic E18 rat brains, as previously described (McCarthy et al. 1980, Frega et al. 2017, Boukas et al. 2019, Fahrner et al. 2019). Animal experiments were conducted in conformity with the Animal Care Committee of the Radboud University Nijmegen Medical Centre, The Netherlands, and conform to the guidelines of the Dutch Council for Animal Care and the European Communities Council Directive 2010/63/EU.

## Acknowledgements

This work was supported by grants from: the Netherlands Organization for Scientific Research, NWO-CAS grant 012.200.001 (to N.N.K); the Netherlands Organization for Health Research and Development ZonMw grant 91217055 (to. H.v.B and N.N.K); SFARI grant 610264 (to N.N.K); ERA-NET NEURON-102 SYNSCHIZ grant (NWO) 013-17-003 4538 (to D.S) and ERA-NET NEURON DECODE! grant (NWO) 013.18.001 (to N.N.K), the Dutch epilepsiefonds WAR 18-02 (to N.N.K.).

## Author contributions

M.F. and N.N.K. conceived and supervised the study. B.M., A.V., E.H., T.K.G., K.L., C.S. and M.F. performed all experiments. B.M., A.V., E.H., G.P. and M.F. performed data analysis. T.K., T.K., H.B., D.S., N.N.K. and M.F. provided resources, conceptualization, and intellectual content. B.M., A.V., E.H. and M.F. wrote the paper. T.K.G., G.P., K.L., C.S., T.K., T.K., H.B., D.S. and N.N.K. edited the paper.

