## Supplementary material for "Human neuronal networks on micro-electrode arrays are a highly robust tool to study disease-specific genotype-phenotype correlations *in vitro*"

### Supplemental material

#### Supplementary figures

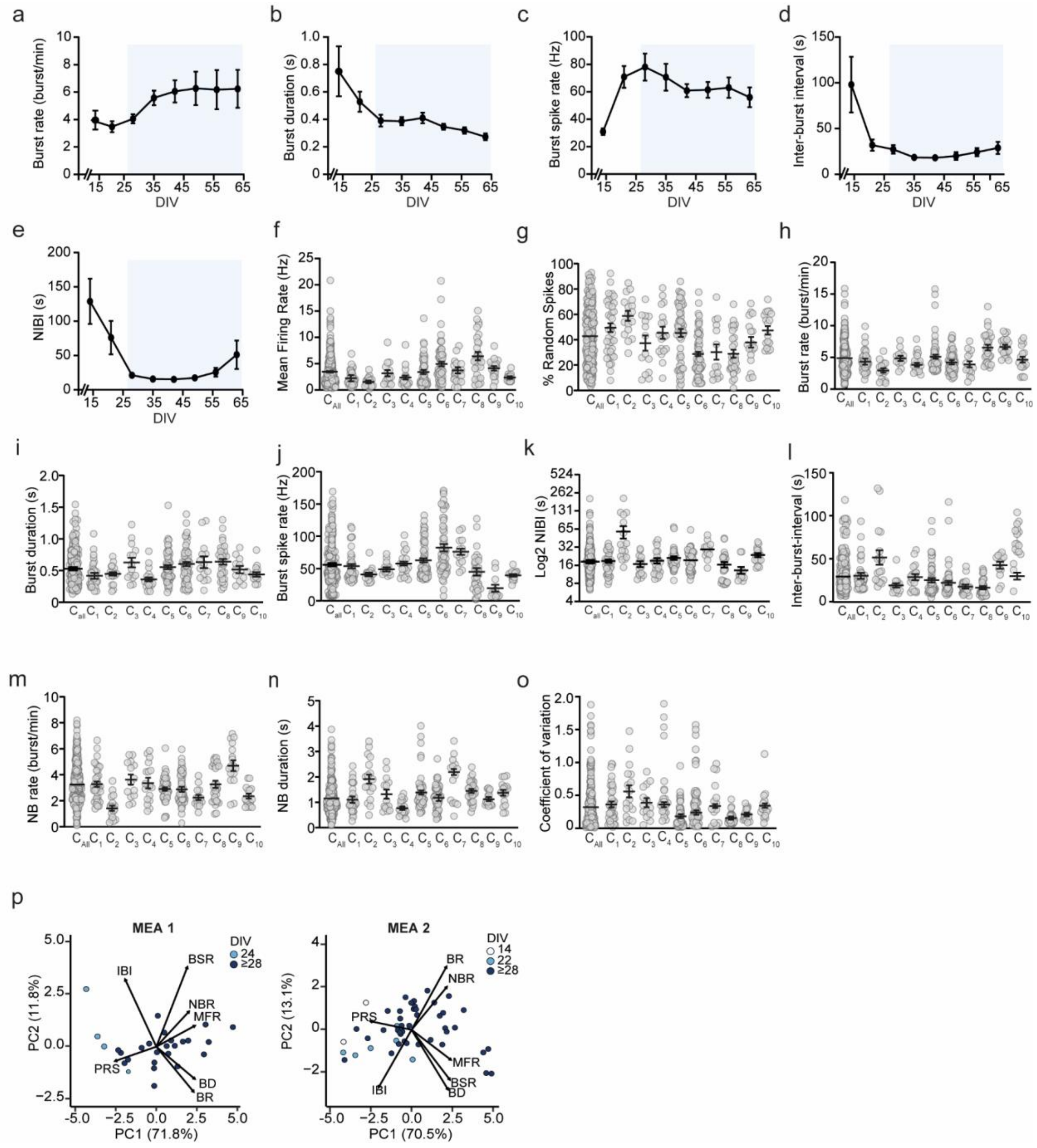

**Figure S1. Variability in neuronal network phenotypes between control lines.** (**a-e**) Neuronal network parameters (representative line C6) develop to reach a certain plateau after DIV 27 (blue box) for (**a**) BR, (**b**) BD, (**c**) BSR, (**d**) IBI and (**e**) NIBI. (**f-o**) Comparison of the MEA parameters (**f**) MFR, (**g**) PRS, (**h**) BR, (**i**) BD, (**k**) BSR, (**l**) NIBI (on log2 scale), (**m**) IBI, (**n**) NBR, (**o**) NBD, (**p**) $CV_{NIBI}$ between all 10 control lines. Kruskal Wallis Anova with Dunn’s correction for multiple testing was used to compare between control lines (**Table S3**). (**q**) PCA plots on all parameters showing data of one control line ($C_6$) pooled and color-coded by DIV (two independent MEA plates are analyzed). DIV = days in vitro, BR = mean burst rate, BD = mean burst duration, BSR = burst spike rate, IBI = inter-burst interval, NIBI = Network burst IBI, MFR = mean firing rate, PRS = percentage of random spike, NBR = network burst rate, NBD = network burst duration, $CV_{NIBI}$ = coefficient of variation on NIBI.

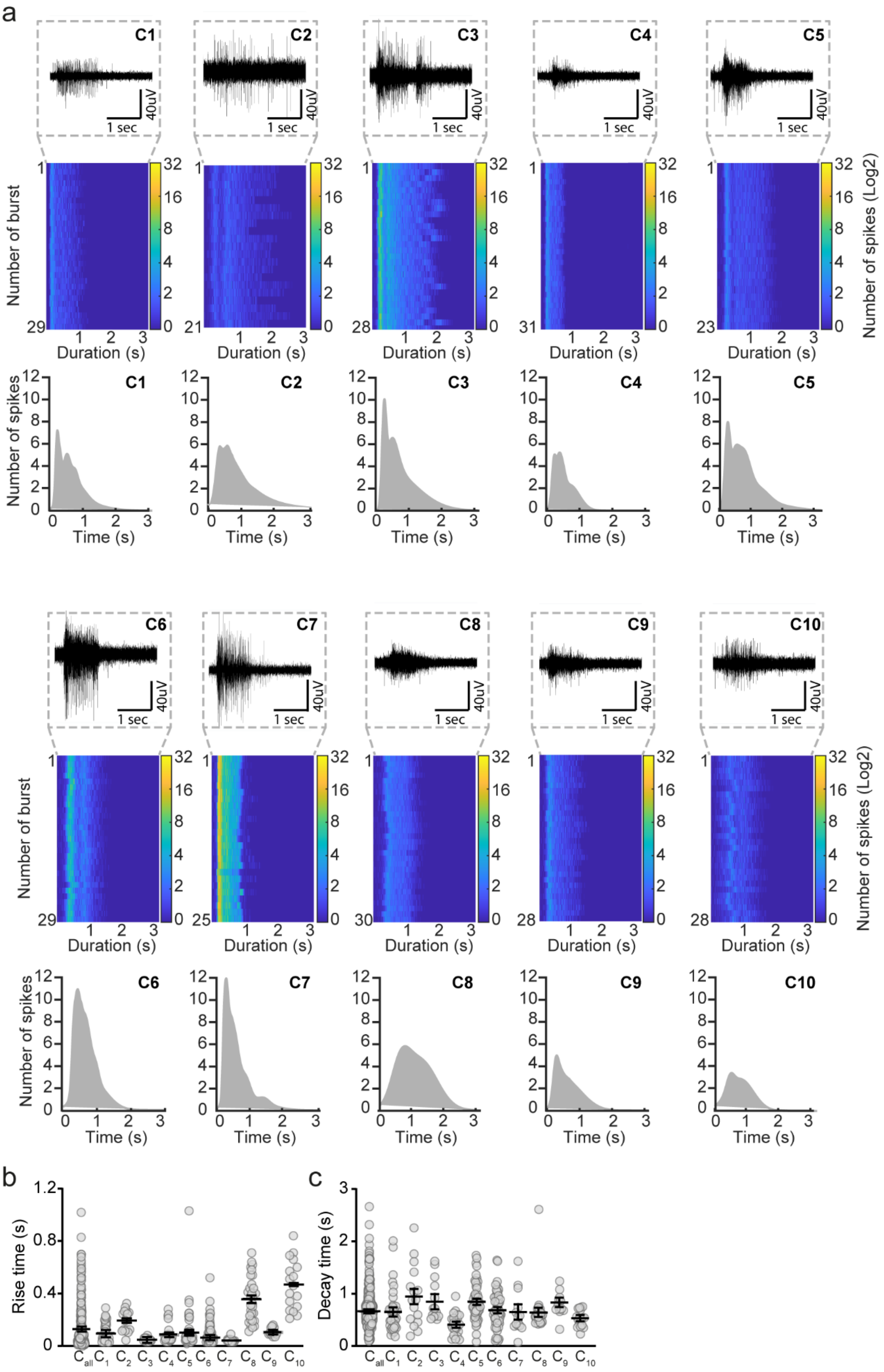
a
C1
C2
C3
C4
C5
C6
C7
C8
C9
C10
1 sec
40uV
Number of burst
Duration (s)
Number of spikes (Log2)
Number of spikes
Time (s)
b
Rise time (s)
c
Decay time (s)
$C_{all}$ $C_1$ $C_2$ $C_3$ $C_4$ $C_5$ $C_6$ $C_7$ $C_8$ $C_9$ $C_{10}$

**Figure S2. Average burst shapes of control lines**. (**a**) Representative network burst trace (zoom in) and alignment from one recording of control lines. Bottom panel: average burst shape from all recordings (Sample size n for $C_1$ = 38, $C_2$ = 15, $C_3$ = 14, $C_4$ = 23, $C_5$ = 55, $C_6$ = 58, $C_7$ = 14, $C_8$ = 30, $C_9$ = 12 and $C_{10}$ = 17). Average burst shapes were used to calculate the (**b**) RT and (**c**) DT. Kruskal Wallis Anova with Dunn's correction for multiple testing was used to compare between control lines (**Table S5**). RT = Rise time, DT = decay time.

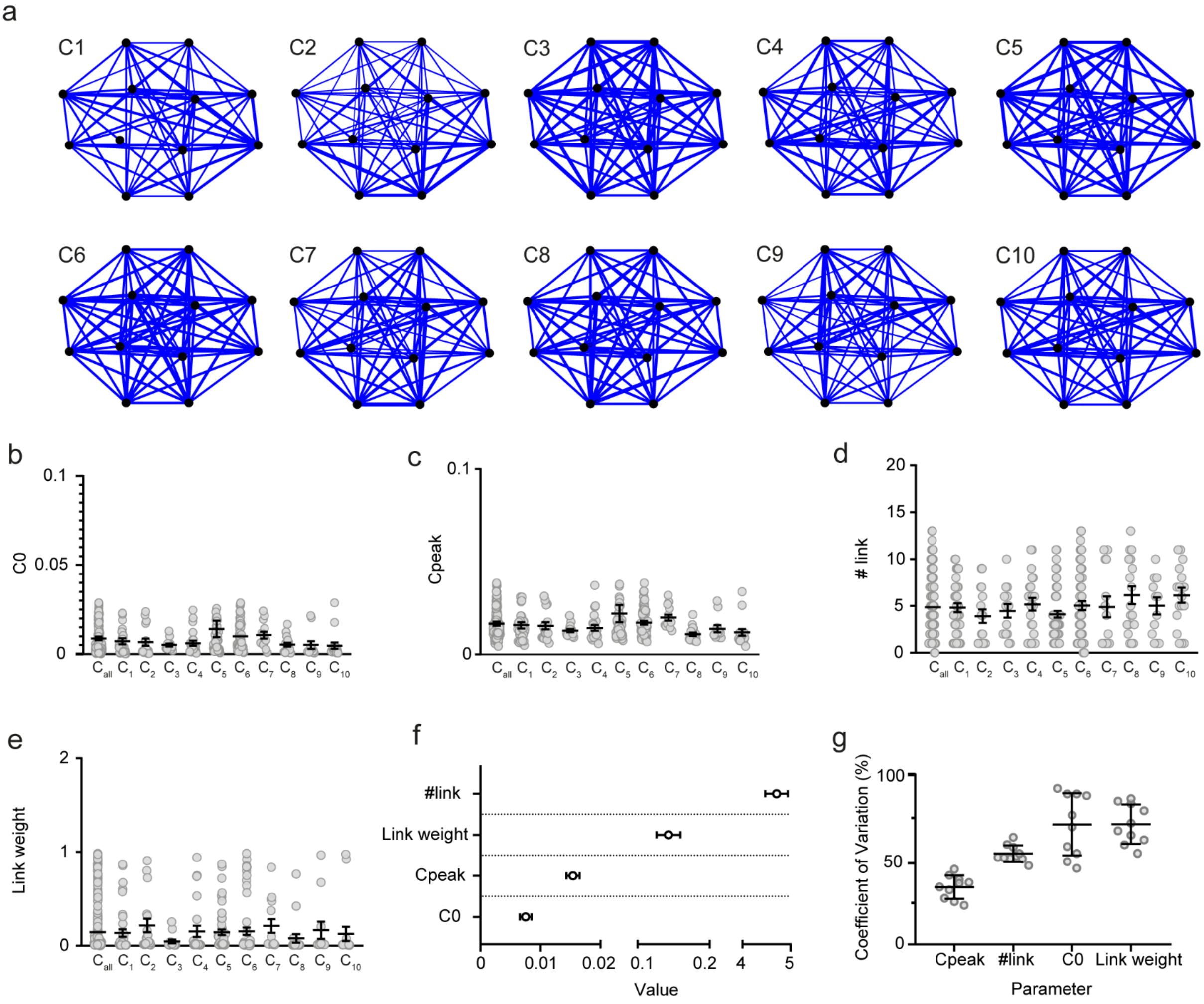

**Figure S3**. **Connectivity in control neuronal networks.** (**a**) Representative connectivity maps showing the links (blue lines) and their strengths (magnitude of the blue lines) (**b-e**) Comparison of the MEA parameters (**b**) $C_0$, (**c**) $C_{peak}$, (**d**) number of link and (**e**) weight of links between all 10 control lines. (**f**) Graph showing the range in which MEA parameters $C_0$, $C_{peak}$, link weight and number of links of all 10 control lines behave (mean ± 95% confidence interval). Values are first averaged per control line, and then averaged across all control lines. (**g**) Percent coefficient of variation explaining the stability of the respective MEA parameter across all 10 control lines (mean ± standard deviation of the mean). N = 278 wells. Kruskal Wallis Anova with Dunn's correction for multiple testing was used to compare between control lines (**Table S2**).

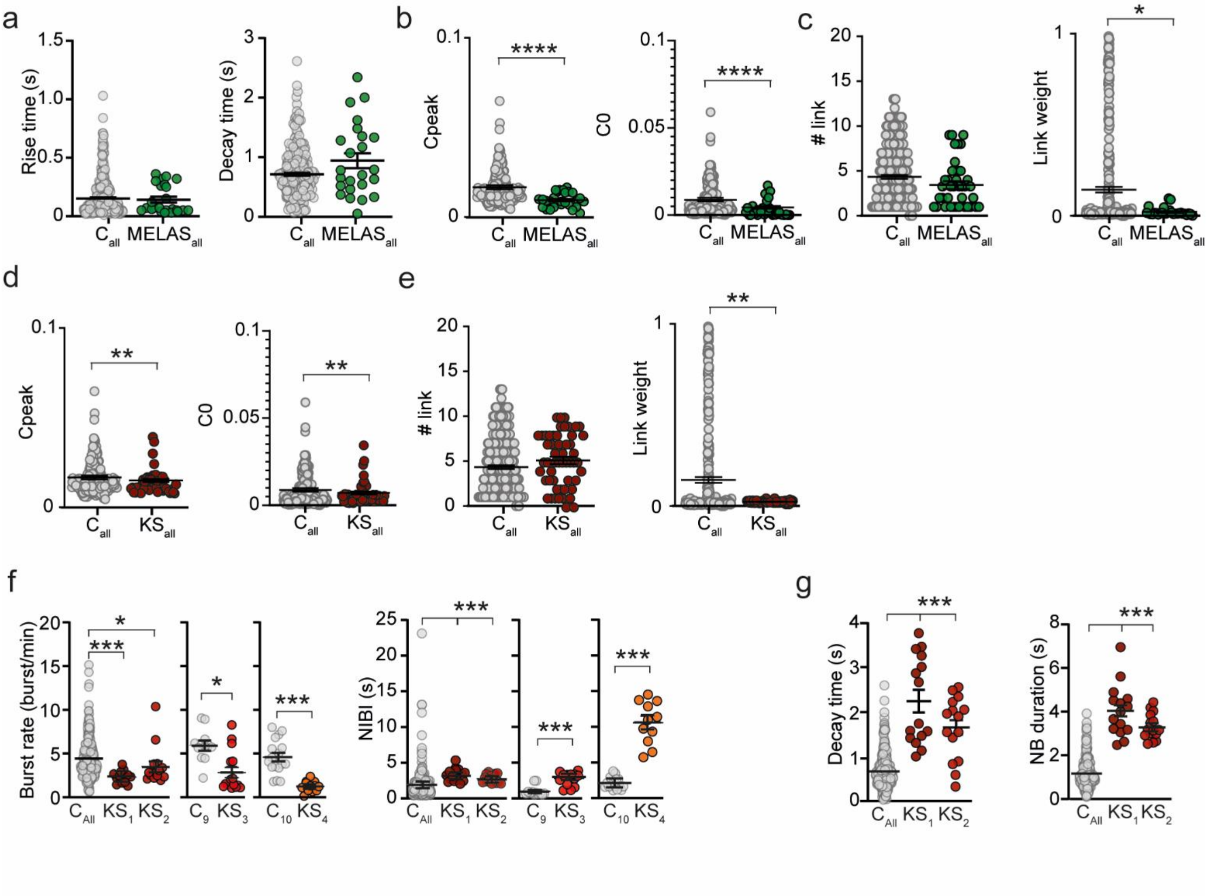

**Figure S4**. **Additional parameters affected in MELAS and KS.** (**a-c**) Comparison of MEA parameters (**a**) RT and DT, (**b**) $C_{peak}$, $C_0$ and (**c**) link number and link weight for MELAS samples versus all controls (Mann Whitney U test with Bonferroni correction for multiple testing was used to compare between patient lines and controls). (**d-e**) Comparison of MEA parameters (**d**) $C_{peak}$, $C_0$ and (**e**) link number and link weight for KS samples versus all controls (Mann Whitney U test with Bonferroni correction for multiple testing was used to compare between patient lines and controls). (**f-g**) Comparison of MEA parameters (**f**) BR and NIBI for all $KS_1$ and $KS_2$ compared to all controls, and for $KS_3$ and $KS_4$ compared to their isogenic controls and (**g**) DT and NBD for $KS_1$ and $KS_2$ compared to all controls (Kruskal Wallis Anova with Dunn's correction for multiple testing was used to compare between control lines). (**Table S8**). $p = 0.05$ *, $p = 0.01$ **, $p = 0.001$ *** and $p<0.0001$ ****. DIV = days in vitro, BR = mean burst rate, NIBI = Network burst inter-burst-interval, RT = Rise time, DT = decay time, BD = burst duration, NBR = network burst rate, NBD = network burst duration. All means, *p*-values and used statistic tests are reported in supplementary Table S8.

**Supplementary tables**

| *Parameter* | *Short name* | *Unit* | *Explanation* |
|---|---|---|---|
| *Mean firing rate* | MFR | Spikes/s | The MFR is a measure of all spikes detected in each electrode during time, which is averaged for all 12 electrodes in a well. |
| *Percentage of random spikes* | PRS | % | The PRS represents the percentage of all spikes that are not organized into a random burst, or a network burst. |
| *Mean burst rate* | MBR | Burst/ minute | A burst is detected when at least 4 spikes are 50 ms or less spaced from each other. The MBR is a measure of all burst detected in each channel in time, which is averaged for all 12 electrodes in a well. |
| *Mean burst duration* | MBD | s | The MBD represents the duration of the burst. |
| *Inter-burst interval* | IBI | s | The IBI the interval between two consecutive bursts. |
| *Burst spike rate* | BSR | Spikes/s | The BSR is the number of spikes detected in a burst divided by the duration of the burst. |
| *Network burst rate* | NBR | Networ k burst/ minute | When a burst is occurring in more than 6 channels at the same time, and 6 are time locked, these bursts are classified as network burst. |
| *Network burst duration* | NBD | s | The NBD represents the duration of the network burst. |
| *Network inter-burst interval* | NIBI | s | The NIBI is the interval between two consecutive network bursts. |
| *Coefficient of variation on NIBI* | $CV_{NIBI}$ | | The coefficient of variation is calculated by dividing the standard deviation of all NIBIs to the mean. The value ranges between 0 (very regular network burst) to 1 (very irregular network burst). |
| *Average burst shape* | | | The average burst shape is an averaged histogram representing the number of spikes occurring into all network burst detected in one well. |
| *Rise time* | RT | s | The rise time is extracted by the average burst shape histogram by fitting it with a Gaussian and calculating the slope between the 20 and 80% of the peak on the rising edge. |
| *Decay time* | DT | s | The decay time is extracted by the average burst shape histogram by fitting it with a Gaussian and calculating the slope between 80 and 20% of the peak on the falling edge. |
| $C_0$ | | | $C_0$ represent the value of the cross correlogram function in the central time bin and indicates the degree of synchronization between two electrodes. |
| $C_{peak}$ | | | $C_{peak}$ represent the peak value of the cross correlogram and indicates the degree of correlation between two electrodes. |
| *Number of links* | | | The number of links indicates the total number of connections between all MEAs electrodes. |
| *Link weight* | | | The link weight indicates the averaged strength of all links detected between all MEAs electrodes |

**Supplementary table 1:** Explanation of extracted MEA parameters.

| *Panel* | | *MFR* | *PRS* | *MBR* | *MBD* | *BSR* | *IBI* | *NBR* | *NBD* | *NIBI* | *$CV_{NIBI}$* | *RT* | *DT* |
|---|---|---|---|---|---|---|---|---|---|---|---|---|---|
| *i* | Mean | 3,5 | 41,3 | 4,8 | 0,51 | 58,5 | 27,47 | 3,2 | 1,28 | 22,68 | 0,3 | 0,15 | 0,71 |
| | SEM | 0,2 | 1,3 | 0,1 | 0,02 | 1,8 | 1,20 | 0,1 | 0,04 | 1,16 | 0,0 | 0,01 | 0,02 |
| | Min | 0,4 | 3,0 | 1,0 | 0,12 | 8,5 | 5,08 | 0,3 | 0,19 | 7,85 | 0,0 | 0,01 | 0,07 |
| | Max | 20,9 | 92,3 | 15,5 | 1,53 | 170,2 | 131,99 | 7,0 | 4,02 | 222,80 | 1,9 | 1,03 | 2,61 |
| *j* | Mean | 62,96 | 46,37 | 39,38 | 42,88 | 39,63 | 58,21 | 38,37 | 40,60 | 43,79 | 74,77 | 58.49 | 48,57 |
| | STD | 17,39 | 14,53 | 9,337 | 7,749 | 20,14 | 13,34 | 9,218 | 10,56 | 9,092 | 18,73 | 26,93 | 12,46 |
| | Min | 29,38 | 26,40 | 26,37 | 28,63 | 17,73 | 33,71 | 28,25 | 20,10 | 30,54 | 47,21 | 28,61 | 29,27 |
| | Max | 87,68 | 73,74 | 57,78 | 51,33 | 84,03 | 80,75 | 61,30 | 54,24 | 93,80 | 103,7 | 141,3 | 64,41 |

| | | *$C_{peak}$* | *$C_0$* | *# link* | *Link weight* |
|---|---|---|---|---|---|
| *f* | Mean | 0,007 | 0,004 | 4,712 | 0,143 |
| | SEM | 0,001 | 0,0009 | 0,234 | 0,016 |
| | Min | 0,010 | 0,004 | 3,647 | 0,045 |
| | Max | 0,022 | 0,014 | 5,889 | 0,216 |
| *g* | Mean | 46,79 | 75,3 | 60,86 | 76,1 |
| | STD | 13,75 | 26,57 | 9,736 | 13,13 |
| | Min | 25,45 | 48,76 | 56,98 | 63,7 |
| | Max | 67,95 | 92,5 | 75,05 | 85,1 |

**Supplementary table 2:** Values of MEA parameters presented **in Figure 1i, j** and **Figure S3f, g. Panel i, f:** Range of each MEA parameters in which controls behave**. Panel j, g:** Percentage of variation of each MEA parameter**.**

| *Panel* | *Dunn's multiple comparisons test* | *n* | *Mean 1* | *SEM* | *n* | *Mean 2* | *SEM* | *Adjusted P-Value* |
|---|---|---|---|---|---|---|---|---|
| *g* | C1 vs. C8 | 38 | 2,278 | 0,222 | 30 | 6,294 | 0,743 | 0,0002 |
| | C1 vs. C6 | 38 | 2,278 | 0,222 | 58 | 4,966 | 0,504 | 0,0014 |
| | C2 vs. C8 | 17 | 1,761 | 0,218 | 30 | 6,294 | 0,743 | 0,0002 |
| | C2 vs. C6 | 17 | 1,761 | 0,218 | 58 | 4,966 | 0,504 | 0,0015 |
| | C4 vs. C8 | 23 | 2,026 | 0,371 | 30 | 6,294 | 0,743 | <0,0001 |
| | C4 vs. C6 | 23 | 2,026 | 0,371 | 58 | 4,966 | 0,504 | 0,0002 |
| | C8 vs. $C_{all}$ | 30 | 6,294 | 0,743 | 278 | 3,477 | 0,171 | 0,006 |
| | $C_{all}$ vs. C6 | 278 | 3,477 | 0,171 | 58 | 4,966 | 0,504 | 0,0346 |
| *h* | C1 vs. C6 | 38 | 49,45 | 3,487 | 58 | 28,56 | 2,302 | 0,0002 |
| | C2 vs. C7 | 17 | 58,9 | 3,771 | 14 | 30,35 | 5,981 | 0,0131 |
| | C2 vs. C8 | 17 | 58,9 | 3,771 | 30 | 32,74 | 3,047 | 0,0044 |
| | C2 vs. C9 | 17 | 58,9 | 3,771 | 12 | 31,91 | 3,329 | 0,0486 |
| | C2 vs. C6 | 17 | 58,9 | 3,771 | 58 | 28,56 | 2,302 | <0,0001 |
| | C5 vs. C6 | 52 | 45,03 | 3,165 | 58 | 28,56 | 2,302 | 0,0031 |
| | C4 vs. C6 | 23 | 49,79 | 4,083 | 58 | 28,56 | 2,302 | 0,0025 |
| | C10 vs. C6 | 17 | 47,36 | 3,317 | 58 | 28,56 | 2,302 | 0,0419 |
| | $C_{all}$ vs. C6 | 278 | 42,34 | 1,303 | 58 | 28,56 | 2,302 | 0,0005 |

| | | | | | | | | |
|---|---|---|---|---|---|---|---|---|
| *i* | C1 vs. C8 | 38 | 4,377 | 0,274 | 30 | 6,895 | 0,432 | *0,0007* |
| | C2 vs. C8 | 17 | 2,928 | 0,344 | 30 | 6,895 | 0,432 | *<0,0001* |
| | C2 vs. C9 | 17 | 2,928 | 0,344 | 12 | 6,304 | 0,533 | *0,0002* |
| | C2 vs. $C_{all}$ | 17 | 2,928 | 0,344 | 278 | 4,812 | 0,133 | *0,0129* |
| | C5 vs. C8 | 52 | 4,863 | 0,389 | 30 | 6,895 | 0,432 | *0,0004* |
| | C4 vs. C8 | 23 | 3,929 | 0,299 | 30 | 6,895 | 0,432 | *<0,0001* |
| | C4 vs. C9 | 23 | 3,929 | 0,299 | 12 | 6,304 | 0,533 | *0,0208* |
| | C7 vs. C8 | 14 | 4,215 | 0,521 | 30 | 6,895 | 0,432 | *0,0152* |
| | C8 vs. $C_{all}$ | 30 | 6,895 | 0,432 | 278 | 4,812 | 0,133 | *0,0001* |
| | C8 vs. C6 | 30 | 6,895 | 0,432 | 58 | 4,565 | 0,220 | *0,0011* |
| *j* | *C1 vs. C8* | 38 | 0,420 | 0,031 | 30 | 0,636 | 0,051 | 0,0285 |
| | *C3 vs. C4* | 14 | 0,627 | 0,078 | 23 | 0,338 | 0,032 | 0,0211 |
| | *C5 vs. C4* | 52 | 0,557 | 0,036 | 23 | 0,338 | 0,032 | 0,00336 |
| | *C4 vs. C7* | 23 | 0,338 | 0,032 | 14 | 0,637 | 0,087 | 0,0204 |
| | *C4 vs. C8* | 23 | 0,338 | 0,032 | 30 | 0,636 | 0,051 | 0,000234 |
| | *C4 vs.* $C_{all}$ | 23 | 0,338 | 0,032 | 278 | 0,514 | 0,015 | 0,00978 |
| | *C4 vs. C6* | 23 | 0,338 | 0,032 | 58 | 0,593 | 0,037 | 0,000328 |
| *k* | *C1 vs. C9* | 38 | 54,83 | 3,906 | 12 | 21,29 | 5,165 | 0,045343 |
| | *C2 vs. C6* | 38 | 54,83 | 3,906 | 58 | 83,59 | 5,663 | 0,007459 |
| | *C2 vs. C5* | 17 | 40,56 | 2,647 | 52 | 64,63 | 3,876 | 0,026287 |
| | *C2 vs. C4* | 17 | 40,56 | 2,647 | 23 | 57,74 | 3,625 | 0,000573 |
| | *C2 vs. C6* | 17 | 40,56 | 2,647 | 58 | 83,59 | 5,663 | 0,000026 |
| | *C5 vs. C9* | 52 | 64,63 | 3,876 | 12 | 21,29 | 5,165 | 0,000179 |
| | *C4 vs. C9* | 23 | 57,74 | 3,625 | 12 | 21,29 | 5,165 | 0,003922 |
| | *C7 vs. C8* | 14 | 77,19 | 5,531 | 30 | 49,13 | 5,441 | 0,017745 |
| | *C7 vs. C9* | 14 | 77,19 | 5,531 | 12 | 21,29 | 5,165 | 0,000005 |
| | *C7 vs. C10* | 14 | 77,19 | 5,531 | 17 | 40,95 | 1,761 | 0,001187 |
| | *C8 vs. C6* | 30 | 49,13 | 5,441 | 58 | 83,59 | 5,663 | 0,001037 |
| | *C9 vs.* $C_{all}$ | 12 | 21,29 | 5,165 | 278 | 58,47 | 1,795 | 0,000956 |
| | *C9 vs. C6* | 12 | 21,29 | 5,165 | 58 | 83,59 | 5,663 | <0,000001 |
| | *C10 vs. C6* | 17 | 40,95 | 1,761 | 58 | 83,59 | 5,663 | 0,000077 |
| | $C_{all}$ *vs. C6* | 278 | 58,47 | 1,795 | 58 | 83,59 | 5,663 | 0,000557 |
| *l* | *C1 vs. C2* | 38 | 19,197 | 1,251 | 16 | 71,407 | 19,425 | 0,001025 |
| | *C1 vs. C7* | 38 | 19,197 | 1,251 | 14 | 30,644 | 2,501 | 0,010311 |
| | *C2 vs. C3* | 16 | 61,109 | 57,32 | 14 | 17,177 | 1,904 | 0,001741 |
| | *C2 vs. C5* | 16 | 61,109 | 57,32 | 52 | 21,605 | 1,467 | 0,015244 |
| | *C2 vs. C4* | 16 | 61,109 | 57,32 | 23 | 19,481 | 1,958 | 0,002403 |
| | *C2 vs. C8* | 16 | 61,109 | 57,32 | 30 | 16,761 | 1,784 | 0,000003 |
| | *C2 vs. C9* | 16 | 61,109 | 57,32 | 12 | 13,471 | 1,901 | 0,000003 |
| | *C2 vs.* $C_{all}$ | 16 | 61,109 | 57,32 | 278 | 22,675 | 1,158 | 0,00043 |
| | *C2 vs. C6* | 16 | 61,109 | 57,32 | 57 | 21,137 | 1,153 | 0,012846 |
| | *C3 vs. C7* | 14 | 17,177 | 1,904 | 14 | 30,644 | 2,501 | 0,009402 |
| | *C4 vs. C7* | 23 | 19,481 | 1,958 | 14 | 30,644 | 2,501 | 0,016409 |
| | *C7 vs. C8* | 14 | 30,644 | 2,501 | 30 | 16,761 | 1,784 | 0,000066 |

| | | | | | | | |
|---|---|---|---|---|---|---|---|
| | *C7 vs. C9* | 14 | 30,644 | 2,501 | 12 | 13,471 | 1,901 | 0,000029 |
| | *C7 vs.* $C_{all}$ | 14 | 30,644 | 2,501 | 278 | 22,675 | 1,158 | 0,008605 |
| | *C8 vs. C10* | 30 | 16,761 | 1,784 | 17 | 24,252 | 1,801 | 0,026793 |
| | *C9 vs. C10* | 12 | 13,471 | 1,901 | 17 | 24,252 | 1,801 | 0,00596 |
| *m* | C1 vs. C8 | 38 | 30,43 | 2,816 | 30 | 16,80 | 1,816 | 0,0007 |
| | C1 vs. C6 | 38 | 30,43 | 2,816 | 58 | 21,95 | 2,360 | 0,031 |
| | C2 vs. C3 | 15 | 52,56 | 7,200 | 14 | 19,18 | 2,636 | 0,0093 |
| | C2 vs. C5 | 17 | 51,29 | 8,637 | 52 | 25,71 | 2,568 | 0,0138 |
| | C2 vs. C7 | 17 | 51,29 | 8,637 | 14 | 17,65 | 2,676 | 0,0016 |
| | C2 vs. C8 | 17 | 51,29 | 8,637 | 30 | 16,80 | 1,816 | <0,0001 |
| | C2 vs. $C_{all}$ | 17 | 51,29 | 8,637 | 273 | 27,47 | 1,196 | 0,0154 |
| | C2 vs. C6 | 17 | 51,29 | 8,637 | 58 | 21,95 | 2,360 | 0,0003 |
| | C3 vs. C9 | 14 | 19,18 | 2,636 | 12 | 41,85 | 4,072 | 0,0203 |
| | C7 vs. C9 | 14 | 17,65 | 2,676 | 12 | 41,85 | 4,072 | 0,0481 |
| | C8 vs. C9 | 30 | 16,80 | 1,816 | 12 | 41,85 | 4,072 | 0,0044 |
| | C8 vs. C10 | 30 | 16,80 | 1,816 | 17 | 30,74 | 3,606 | <0,0001 |
| | C8 vs. $C_{all}$ | 30 | 16,80 | 1,816 | 273 | 27,47 | 1,196 | 0,0363 |
| | C9 vs. C6 | 12 | 41,85 | 4,072 | 58 | 21,95 | 2,360 | 0,0162 |
| *n* | C1 vs. C2 | 38 | 3,363 | 0,208 | 17 | 1,465 | 0,218 | <0,0001 |
| | C2 vs. C3 | 17 | 1,465 | 0,218 | 14 | 3,736 | 0,366 | 0,0003 |
| | C2 vs. C5 | 17 | 1,465 | 0,218 | 52 | 2,902 | 0,125 | 0,0014 |
| | C2 vs. C4 | 17 | 1,465 | 0,218 | 23 | 3,465 | 0,278 | 0,0001 |
| | C2 vs. C8 | 17 | 1,465 | 0,218 | 30 | 4,067 | 0,314 | <0,0001 |
| | C2 vs. C9 | 17 | 1,465 | 0,218 | 12 | 4,642 | 0,379 | <0,0001 |
| | C2 vs. $C_{all}$ | 17 | 1,465 | 0,218 | 273 | 3,177 | 0,084 | <0,0001 |
| | C2 vs. C6 | 17 | 1,465 | 0,218 | 57 | 2,928 | 0,144 | 0,004 |
| | C7 vs. C8 | 14 | 2,229 | 0,203 | 30 | 4,067 | 0,314 | 0,0036 |
| | C7 vs. C9 | 14 | 2,229 | 0,203 | 12 | 4,642 | 0,379 | 0,0007 |
| | C8 vs. C10 | 30 | 4,067 | 0,314 | 17 | 2,306 | 0,175 | 0,0029 |
| | C9 vs. C10 | 12 | 4,642 | 0,379 | 17 | 2,306 | 0,175 | 0,0006 |
| | C9 vs. C6 | 12 | 4,642 | 0,379 | 57 | 2,928 | 0,144 | 0,0273 |
| *o* | *C1 vs. C2* | 38 | 1,094 | 0,079 | 17 | 1,887 | 0,205 | 0,0124 |
| | *C1 vs. C8* | 38 | 1,094 | 0,079 | 30 | 1,498 | 0,076 | 0,0265 |
| | *C2 vs. C4* | 17 | 1,887 | 0,205 | 23 | 0,704 | 0,060 | <0,0001 |
| | *C5 vs. C4* | 52 | 1,379 | 0,100 | 23 | 0,704 | 0,060 | 0,0002 |
| | *C4 vs. C7* | 23 | 0,704 | 0,060 | 14 | 1,925 | 0,239 | <0,0001 |
| | *C4 vs. C8* | 23 | 0,704 | 0,060 | 30 | 1,498 | 0,076 | <0,0001 |
| | *C4 vs. C10* | 23 | 0,704 | 0,060 | 17 | 1,375 | 0,113 | 0,0011 |
| | *C4 vs.* $C_{all}$ | 23 | 0,704 | 0,060 | 273 | 1,277 | 0,038 | <0,0001 |
| | *C4 vs. C6* | 23 | 0,704 | 0,060 | 57 | 1,202 | 0,062 | 0,0013 |
| *p* | *C1 vs. C5* | 38 | 0,364 | 0,045 | 52 | 0,188 | 0,025 | 0,0246 |
| | *C1 vs. C8* | 38 | 0,364 | 0,045 | 30 | 0,154 | 0,017 | 0,048 |
| | *C2 vs. C5* | 17 | 0,549 | 0,098 | 52 | 0,188 | 0,025 | 0,0013 |

| | | | | | | | |
|---|---|---|---|---|---|---|---|
| *C2 vs. C8* | 17 | 0,549 | 0,098 | 30 | 0,154 | 0,017 | 0,0023 |
| *C5 vs. C4* | 52 | 0,188 | 0,025 | 23 | 0,587 | 0,111 | <0,0001 |
| *C5 vs.* $C_{all}$ | 52 | 0,188 | 0,025 | 278 | 0,331 | 0,019 | 0,0181 |
| *C5 vs. C6* | 52 | 0,188 | 0,025 | 55 | 0,380 | 0,051 | 0,0271 |
| *C4 vs. C8* | 23 | 0,587 | 0,111 | 30 | 0,154 | 0,017 | 0,0002 |

**Supplementary table 3:** Statistics of MEA parameters presented **in Supplementary figure 1f-o and Supplementary figure 3b-e.** Only comparisons that are statistically different are reported in the table. No statistical difference was found when comparing the parameters in Supplementary figure 3b-e. Sample size n represents the number of recorded MEA wells between DIV27-35. All data represent means ± SEM. Kruskal Wallis Anova with Dunn's correction for multiple testing was used to compare between control lines. DIV = Days in vitro.

| *Panel* | *Dunn's multiple comparisons test* | *n* | *Mean 1* | *SEM* | *n* | *Mean 2* | *SEM* | *Adjusted P Value* |
|---|---|---|---|---|---|---|---|---|
| *g* | $C_{all}$ vs $C_{suboptimal.}$ | 278 | 3,177 | 0,084 | 48 | 1,632 | 0,306 | <0,0001 |
| | $C_{optimal}$ vs $C_{suboptimal.}$ | 48 | 3,673 | 0,213 | 48 | 1,632 | 0,306 | <0,0001 |
| | $C_{all}$ vs $C_{suboptimal.}$ | 278 | 22,675 | 1,158 | 48 | 302,649 | 41,648 | <0,0001 |
| | $C_{optimal}$ vs $C_{suboptimal.}$ | 48 | 18,940 | 1,693 | 48 | 302,649 | 41,648 | <0,0001 |
| | $C_{all}$ vs $C_{suboptimal.}$ | 278 | 0,340 | 0,020 | 48 | 0,690 | 0,060 | <0,0001 |
| | $C_{optimal}$ vs $C_{suboptimal.}$ | 48 | 0,444 | 0,691 | 48 | 0,690 | 0,060 | 0,00062 |
| *h* | $C_1$ vs $C_{active\ el.}$ | 27 | 2,257 | 0,232 | 14 | 1,876 | 0,253 | ns |
| | $C_1$ vs $C_{all\ el.}$ | 27 | 2,257 | 0,232 | 14 | 0,771 | 0,098 | <0,0001 |
| | $C_1$ vs $C_{active\ el.}$ | 27 | 3,022 | 0,199 | 14 | 3,493 | 0,727 | ns |
| | $C_1$ vs $C_{all\ el.}$ | 27 | 3,022 | 0,199 | 14 | 0,236 | 0,192 | <0,0001 |

**Supplementary table 4:** Statistics of MEA parameters presented **in Figure 2g, h.** Sample size n represents the number of recorded MEA wells between DIV27-35. Panel f: Kruskal Wallis Anova with Dunn's correction for multiple testing was used to compare between control lines. Panel g: One-way Anova with Tukey correction for multiple testing was used to compare between control lines.

| *Panel* | *Dunn's multiple comparisons test* | *n* | *Mean 1* | *SEM* | *n* | *Mean 2* | *SEM* | *Adjusted P Value* |
|---|---|---|---|---|---|---|---|---|
| *b* | *C1 vs. C8* | 38 | 0,098 | 0,012 | 30 | 0,360 | 0,031 | <0,0001 |
| | *C1 vs. C10* | 38 | 0,098 | 0,012 | 17 | 0,468 | 0,046 | <0,0001 |
| | *C2 vs. C3* | 16 | 0,197 | 0,021 | 14 | 0,049 | 3,759 | 0,0023 |
| | *C2 vs. C5* | 16 | 0,197 | 0,021 | 55 | 0,105 | 0,020 | 0,0319 |
| | *C2 vs. C7* | 16 | 0,197 | 0,021 | 14 | 0,043 | 4,118 | 0,0001 |
| | *C3 vs. C8* | 14 | 0,049 | 3,759 | 30 | 0,360 | 0,031 | <0,0001 |
| | *C3 vs. C10* | 14 | 0,049 | 3,759 | 17 | 0,468 | 0,046 | <0,0001 |
| | *C5 vs. C8* | 55 | 0,105 | 0,020 | 30 | 0,360 | 0,031 | <0,0001 |
| | *C5 vs. C10* | 55 | 0,105 | 0,020 | 17 | 0,468 | 0,046 | <0,0001 |
| | *C4 vs. C8* | 23 | 0,088 | 0,016 | 30 | 0,360 | 0,031 | <0,0001 |
| | *C4 vs. C10* | 23 | 0,088 | 0,016 | 17 | 0,468 | 0,046 | <0,0001 |
| | *C7 vs. C8* | 14 | 0,043 | 4,118 | 30 | 0,360 | 0,031 | <0,0001 |
| | *C7 vs. C9* | 14 | 0,043 | 4,118 | 12 | 0,108 | 8,947 | 0,0473 |
| | *C7 vs. C10* | 14 | 0,043 | 4,118 | 17 | 0,468 | 0,046 | <0,0001 |
| | *C7 vs.* $C_{all}$ | 14 | 0,043 | 4,118 | 277 | 0,151 | 9,823 | 0,0079 |
| | *C8 vs.* $C_{all}$ | 30 | 0,360 | 0,031 | 277 | 0,151 | 9,823 | <0,0001 |
| | *C8 vs. C6* | 30 | 0,360 | 0,031 | 58 | 0,100 | 0,012 | <0,0001 |

| | | | | | | | | |
|---|---|---|---|---|---|---|---|---|
| | *C10 vs.* $C_{all}$ | 17 | 0,468 | 0,046 | 277 | 0,151 | 9,823 | <0,0001 |
| | *C10 vs. C6* | 17 | 0,468 | 0,046 | 58 | 0,100 | 0,012 | <0,0001 |

| | | | | | | | | |
|---|---|---|---|---|---|---|---|---|
| *c* | *C1 vs. C2* | 38 | 0,656 | 0,068 | 16 | 0,954 | 0,146 | 0,0236 |
| | *C1 vs. C5* | 38 | 0,656 | 0,068 | 55 | 0,849 | 0,047 | 0,0222 |
| | *C2 vs. C4* | 16 | 0,954 | 0,146 | 23 | 0,413 | 0,042 | <0,0001 |
| | *C2 vs. C8* | 16 | 0,954 | 0,146 | 30 | 0,642 | 0,070 | 0,0407 |
| | *C2 vs. C10* | 16 | 0,954 | 0,146 | 17 | 0,525 | 0,039 | 0,0097 |
| | *C3 vs. C4* | 14 | 0,856 | 0,105 | 23 | 0,413 | 0,042 | 0,0067 |
| | *C5 vs. C4* | 55 | 0,849 | 0,047 | 23 | 0,413 | 0,042 | <0,0001 |
| | *C5 vs. C10* | 55 | 0,849 | 0,047 | 17 | 0,525 | 0,039 | 0,0161 |
| | *C5 vs.* $C_{all}$ | 55 | 0,849 | 0,047 | 277 | 0,703 | 0,023 | 0,0268 |
| | *C4 vs. C9* | 23 | 0,413 | 0,042 | 12 | 0,794 | 0,067 | 0,0036 |
| | *C4 vs.* $C_{all}$ | 23 | 0,413 | 0,042 | 277 | 0,703 | 0,023 | 0,0016 |
| | *C4 vs. C6* | 23 | 0,413 | 0,042 | 58 | 0,686 | 0,045 | 0,0071 |

**Supplementary table 5:** Statistics of MEA parameters presented **in Supplementary figure 2b-c.** Sample size n represents the number of recorded MEA wells between DIV 27-35. All data represent means ± SEM. Kruskal Wallis Anova with Dunn's correction for multiple testing was used to compare between control lines. DIV = Days in vitro.

| *Parameter* | *n* | *Mean* $C_{all}$ | *SEM* | *n* | *Mean* $MELAS_{all}$ | *SEM* | *p-value (DIVs)* |
|---|---|---|---|---|---|---|---|
| *MFR* | 278 | 3,477 | 0,171 | 115 | 0,851 | 0,123 | <0,0001 |
| *PRS* | 278 | 41,290 | 1,268 | 115 | 88,430 | 1,623 | <0,0001 |
| *BD* | 278 | 0,514 | 0,015 | 115 | 0,238 | 0,019 | <0,0001 |
| *NBR* | 278 | 3,177 | 0,084 | 115 | 0,290 | 0,082 | <0,0001 |
| *Parameter* | **n** | **Mean** $C_{all}$ | **SEM** | **n** | **Mean** $KS_{all}$ | **SEM** | **p-value (DIVs)** |
| *BD* | 278 | 0,514 | 0,015 | 58 | 1,363 | 0,103 | <0,0001 |
| *NBR* | 278 | 3,177 | 0,084 | 58 | 1,538 | 0,094 | <0,0001 |
| *NBD* | 278 | 1,277 | 0,038 | 58 | 2,995 | 0,143 | <0,0001 |
| *DT* | 278 | 0,713 | 0,023 | 58 | 1,394 | 0,123 | <0,0001 |

**Supplementary table 6:** Statistics of MEA parameters presented **in Figure 3c-j.** Sample size n represents the number of recorded MEA wells between DIV27-35. All data represent means ± SEM. Mann Whitney U test with Bonferroni correction for multiple testing was used. MFR = Mean firing rate, PRS = Percentage of random spikes, BD = Burst duration, NBR = Network burst rate, NBD = Network burst duration, DT = decay time.

| | *Parameter* | *n* | *Mean 1* | *SEM* | *n* | *Mean 2* | *SEM* | *p-value* |
|---|---|---|---|---|---|---|---|---|
| *C2 vs M1* | MFR | 17 | 1,761 | 0,218 | 24 | 0,792 | 0,180 | 0,00028 |
| | NBR | 17 | 1,465 | 0,218 | 24 | 0,146 | 0,080 | <0,0001 |
| | PRS | 17 | 58,900 | 3,771 | 24 | 92,040 | 1,658 | 8,0E-10 |
| | BD | 17 | 0,458 | 0,037 | 24 | 0,241 | 0,026 | 2,2E-05 |
| *C4 vs M2* | MFR | 23 | 2,026 | 0,371 | 23 | 0,821 | 0,141 | 1,3E-04 |
| | NBR | 23 | 3,465 | 0,278 | 23 | 0,409 | 0,148 | <0,0001 |
| | PRS | 23 | 49,790 | 4,083 | 23 | 86,370 | 3,464 | <0,0001 |
| | BD | 23 | 0,338 | 0,032 | 23 | 0,174 | 0,013 | <0,0001 |
| *C5 vs M3* | MFR | 52 | 3,466 | 0,376 | 68 | 0,881 | 0,193 | <0,0001 |
| | NBR | 52 | 2,902 | 0,125 | 68 | 0,300 | 0,126 | <0,0001 |
| | PRS | 52 | 45,030 | 3,165 | 68 | 87,860 | 2,411 | <0,0001 |

| | | | | | | | | |
|---|---|---|---|---|---|---|---|---|
| | BD | 52 | 0,557 | 0,036 | 68 | 0,259 | 0,030 | <0,0001 |
| *C9 vs KS3* | BD | 12 | 0,535 | 0,055 | 16 | 1,388 | 0,154 | 0,00145 |
| | NBR | 12 | 4,642 | 0,379 | 16 | 1,988 | 0,188 | 0,000004 |
| | NBD | 12 | 1,165 | 0,068 | 16 | 2,642 | 0,185 | 2,63E-07 |
| | DT | 12 | 0,794 | 0,067 | 16 | 0,453 | 0,092 | 0,01164 |
| *C10 vs KS4* | BD | 17 | 0,444 | 0,031 | 12 | 0,740 | 0,051 | 2,51E-07 |
| | MFR | 17 | 2,334 | 0,166 | 12 | 1,103 | 0,094 | 2,69E-07 |
| | NBR | 17 | 2,306 | 0,175 | 12 | 0,500 | 0,048 | 3,85E-08 |
| | DT | 17 | 0,525 | 0,039 | 12 | 1,223 | 0,072 | 3,85E-08 |

**Supplementary table 7:** Statistics of MEA parameters presented **in Figure 4b-e and i-l.** Sample size n represents the number of recorded MEA wells between DIV 28-35. All data represent means ± SEM. Mann Whitney U test with Bonferroni correction for multiple testing was used to compare between patient lines and their isogenic controls. MFR = Mean firing rate, PRS = Percentage of random spikes, BD = Burst duration, NBR = Network burst rate, NBD = Network burst duration, NBR = Network burst duration, DT = decay time. DIV = Days in vitro.

| | *Parameter* | *n* | *Mean 1* | *SEM* | *n* | *Mean 2* | *SEM* | *p-value* |
|---|---|---|---|---|---|---|---|---|
| *$C_{all}$ vs $Melas_{all}$* | RT | 278 | 0,713 | 0,023 | 24 | 0,940 | 0,125 | 0,000007 |
| | DT | 278 | 0,150 | 0,010 | 24 | 0,142 | 0,025 | <0,0001 |
| | C0 | 278 | 0.009 | 0.001 | 24 | 0.003 | 0.0007 | <0,0001 |
| | Cpeak | 278 | 0,016 | 0.001 | 24 | 0.009 | 0.0007 | <0,0001 |
| | # link | 278 | 4.602 | 0.202 | 24 | 3.447 | 0.4245 | 0.0594 |
| | Link weight | 278 | 0.144 | 0.016 | 24 | 0.021 | 0.005 | 0.0047 |
| *$C_{all}$ vs KS1* | BR | 278 | 4,812 | 0,133 | 15 | 2,673 | 0,174 | 0,000007 |
| | NIBI | 278 | 3,177 | 0,084 | 15 | 35,061 | 2,194 | 0,000001 |
| *$C_{all}$ vs KS2* | BR | 278 | 4,812 | 0,133 | 15 | 3,908 | 0,569 | 0,043117 |
| | NIBI | 278 | 3,177 | 0,084 | 15 | 30,360 | 1,441 | 0,000059 |
| *C9 vs KS3* | BR | 12 | 6,304 | 0,533 | 16 | 3,195 | 0,580 | 0,00145 |
| | NIBI | 12 | 4,642 | 0,379 | 16 | 1,988 | 0,188 | 0,000004 |
| *C10 vs KS4* | BR | 17 | 4,946 | 0,4813 | 12 | 1,603 | 0,176 | 2,51E-07 |
| | NIBI | 17 | 2,334 | 0,166 | 12 | 1,103 | 0,094 | 2,69E-07 |
| *$C_{all}$ vs KS1* | BD | 278 | 0,514 | 0,015 | 15 | 2,046 | 0,249 | <0,0001 |
| | NBR | 278 | 3,177 | 0,084 | 15 | 1,600 | 0,097 | <0,0001 |
| | NBD | 278 | 1,277 | 0,383 | 15 | 4,002 | 0,307 | 2E-10 |
| | DT | 278 | 0,703 | 0,023 | 15 | 2,261 | 0,251 | 2,4E-09 |
| *$C_{all}$ vs KS2* | BD | 278 | 0,514 | 0,015 | 15 | 1,151 | 0,127 | <0,0001 |
| | NBR | 278 | 3,177 | 0,084 | 15 | 1,988 | 0,188 | 0,000027 |
| | NBD | 278 | 1,277 | 0,383 | 15 | 3,261 | 0,153 | 1,7E-09 |
| | DT | 278 | 0,703 | 0,023 | 15 | 1,669 | 0,181 | 4,90E-06 |
| *$C_{all}$ vs $KS_{all}$* | C0 | 278 | 0.009 | 0.001 | 58 | 0.005 | 0.0007 | 0.0046 |
| | Cpeak | 278 | 0,016 | 0.001 | 58 | 0.012 | 0.0007 | 0.0033 |
| | # link | 278 | 4.602 | 0.202 | 58 | 5.242 | 0.405 | 0.1033 |
| | Link weight | 278 | 0.144 | 0.016 | 58 | 0.013 | 0.005 | 0.0077 |

**Supplementary table 8:** Statistics of MEA parameters presented **in Supplementary figure 4.** Sample size n represents the number of recorded MEA wells between DIV27-35. All data represent means ± SEM. Mann Whitney U test with Bonferroni correction for multiple testing was used to compare between patient lines and their isogenic controls. Kruskal Wallis Anova with Dunn's correction for multiple testing was used to compare between Call and KS1 and KS2. BD = Burst duration, BR = Burst rate, NBR = Network burst rate, NBD = Network burst duration, NBR = Network burst duration, NIBI = Network burst inter-burst-interval, RT = Rise Time, DT = decay time. DIV = Days in vitro.

| | | *Post hoc power calculation* | | *A priori power calculation* | |
|---|---|---|---|---|---|
| *Compared lines* | **Top parameters** | **# wells used for calculation** | **Power** | **# wells needed for power 0,9** | **Effect size** |
| *C9 vs KS3* | BR | C9 = 12 | 0.999 | 12 | 1.9 |
| | BD | KS3 =16 | 0.999 | 12 | 1.9 |
| | NBR | | 1 | 6 | 3 |
| | NBD | | 0.999 | 8 | 2.7 |
| | NIBI | | 0.999 | 10 | 2.3 |
| *C10 vs KS4* | MFR | C10 = 17 | 0.999 | 12 | 2 |
| | BR | KS4 =12 | 1 | 6 | 3 |
| | NBR | | 0.98 | 14 | 2 |
| | NBD | | 0.97 | 8 | 2.8 |
| | NIBI | | 1 | 6 | 3.4 |
| *C4 versus M2* | MFR | C4 = 23 | 0.999 | 12 | 2 |
| | PRS | M2 = 23 | 1 | 10 | 2.1 |
| | BD | | 0.998 | 20 | 1.4 |
| | BSR | | 0.999 | 16 | 1.5 |
| | NBR | | 1 | 6 | 3 |
| *C5 versus M3* | MFR | C5 = 55 | 1 | 12 | 1.9 |
| | PRS | M3 = 68 | 1 | 12 | 1.9 |
| | BD | | 0.999 | 24 | 1.4 |
| | BRS | | 1 | 22 | 1.3 |
| | NBR | | 1 | 12 | 2 |
| *C2 versus M1* | PRS | C2 = 17 | 1 | 8 | 2.8 |
| | BD | M1 = 24 | 0.999 | 16 | 1.6 |
| | NBR | | 0.927 | 6 | 1 |
| | BSR | | 0.997 | 22 | 1.5 |
| *All controls vs KS* | BR | $C_{all}$ = 278 | 1 | 14 | 2 |
| | BD | $KS_{all}$ = 58 | 1 | 12 | 2.1 |
| | NBR | | 1 | 14 | 2 |
| | NBD | | 1 | 14 | 1.9 |
| | NIBI | | 0.999 | 10 | 2.55 |
| | DT | | 0.999 | 12 | 2.27 |
| *All controls vs MELAS* | MFR | $C_{all}$ = 278 | 1 | 12 | 1.9 |
| | PRS | $MELAS_{all}$ = 115 | 1 | 8 | 2.5 |
| | BD | | 1 | 8 | 2.78 |
| | BSR | | 1 | 10 | 2.3 |
| | NBR | | 1 | 6 | 3 |

**Supplementary table 9:** Power calculations on MEA data. Left: post hoc power calculation on PCA parameters that describe the patient phenotype. Alpha is 0.05. Right: a priori Power calculation to calculate the sample size needed to achieve a power of 0.9 using the PCA parameters that describe the patient phenotype. Effect size represents the magnitude of the difference between populations.

| *Panel* | *Parameter* | *PC1* | *PC2* |
|---|---|---|---|
| *S1q Plate 1* | MFR | 0.15 | 0.07 |
| | PRS | 0.16 | 0.05 |
| | BR | 0.15 | 0.15 |
| | BD | 0.15 | 0.11 |
| | BSR | 0.12 | 0.27 |
| | IBI | 0.12 | 0.23 |
| | NBR | 0.13 | 0.12 |
| *S1q Plate 2* | MFR | 0.15 | 0.10 |
| | PRS | 0.16 | 0.03 |
| | BR | 0.14 | 0.20 |
| | BD | 0.14 | 0.19 |
| | BSR | 0.15 | 0.16 |
| | IBI | 0.13 | 0.18 |
| | NBR | 0.14 | 0.14 |
| *2a-d+h* | MFR | 0.16 | 0.01 |
| | PRS | 0.14 | 0.03 |
| | BR | 0.08 | 0.02 |
| | BD | 0.14 | 0.07 |
| | BSR | 0.06 | 0.06 |
| | IBI | 0.08 | 0.10 |
| | NBR | 0.05 | 0.14 |
| | NBD | 0.08 | 0.16 |
| | NIBI | 0.07 | 0.15 |
| | $CV_{NIBI}$ | 0.07 | 0.04 |
| | RT | 0.01 | 0.08 |
| | DT | 0.05 | 0.14 |
| *2i+j* | MFR | 0.08 | 0.11 |
| | PRS | 0.00 | 0.14 |
| | BR | 0.13 | 0.04 |
| | BD | 0.10 | 0.12 |
| | BSR | 0.09 | 0.09 |
| | IBI | 0.06 | 0.06 |
| | NBR | 0.10 | 0.10 |
| | NBD | 0.11 | 0.08 |
| | NIBI | 0.11 | 0.08 |
| | $CV_{NIBI}$ | 0.06 | 0.01 |
| | RT | 0.04 | 0.11 |
| | DT | 0.11 | 0.06 |
| *3l* | MFR | 0.18 | 0.03 |
| | PRS | 0.19 | 0.08 |

| | | | |
|---|---|---|---|
| | BR | 0.08 | 0.36 |
| | BD | 0.16 | 0.11 |
| | BSR | 0.13 | 0.15 |
| | IBI | 0.11 | 0.21 |
| | NBR | 0.15 | 0.06 |
| *3m* | MFR | 0.08 | 0.15 |
| | PRS | 0.11 | 0.12 |
| | BR | 0.05 | 0.13 |
| | BD | 0.16 | 0.01 |
| | BSR | 0.09 | 0.07 |
| | IBI | 0.01 | 0.12 |
| | NBR | 0.10 | 0.14 |
| | NBD | 0.16 | 0.03 |
| | NIBI | 0.06 | 0.15 |
| | $CV_{NIBI}$ | 0.05 | 0.07 |
| | RT | 0.00 | 0.01 |
| | DT | 0.14 | 0.01 |
| *4g Set 1* | MFR | 0.19 | 0.12 |
| | PRS | 0.20 | 0.09 |
| | BR | 0.05 | 0.32 |
| | BD | 0.18 | 0.15 |
| | BSR | 0.13 | 0.07 |
| | IBI | 0.08 | 0.24 |
| | NBR | 0.17 | 0.03 |
| *4g Set 2* | MFR | 0.16 | 0.04 |
| | PRS | 0.18 | 0.06 |
| | BR | 0.07 | 0.34 |
| | BD | 0.17 | 0.11 |
| | BSR | 0.17 | 0.14 |
| | IBI | 0.11 | 0.26 |
| | NBR | 0.14 | 0.05 |
| *4g Set 3* | MFR | 0.17 | 0.03 |
| | PRS | 0.19 | 0.07 |
| | BR | 0.04 | 0.41 |
| | BD | 0.16 | 0.08 |
| | BSR | 0.17 | 0.09 |
| | IBI | 0.11 | 0.26 |
| | NBR | 0.15 | 0.05 |
| *4n Set 1* | MFR | 0.11 | 0.05 |
| | PRS | 0.04 | 0.11 |
| | BR | 0.12 | 0.01 |
| | BD | 0.12 | 0.00 |

| | | | |
|---|---|---|---|
| | BSR | 0.03 | 0.18 |
| | IBI | 0.01 | 0.19 |
| | NBR | 0.11 | 0.06 |
| | NBD | 0.11 | 0.07 |
| | NIBI | 0.12 | 0.03 |
| | $CV_{NIBI}$ | 0.05 | 0.10 |
| | RT | 0.07 | 0.12 |
| | DT | 0.10 | 0.07 |
| *4n Set 2* | MFR | 0.10 | 0.01 |
| | PRS | 0.04 | 0.22 |
| | BR | 0.10 | 0.00 |
| | BD | 0.08 | 0.12 |
| | BSR | 0.08 | 0.03 |
| | IBI | 0.05 | 0.21 |
| | NBR | 0.11 | 0.06 |
| | NBD | 0.09 | 0.07 |
| | NIBI | 0.11 | 0.01 |
| | $CV_{NIBI}$ | 0.08 | 0.01 |
| | RT | 0.08 | 0.14 |
| | DT | 0.10 | 0.11 |

**Supplementary table 10:** Weight of each parameter on a principal component for all PCA plots. Panels mentioned in the first column correspond to the figure for which data is shown. Parameters used to generate the PCA plot are shown in the second column. Normalized weights are shown per parameter for PC1 and PC2 of each PCA plot. MFR = Mean Firing Rate, PRS = Percentage of random spikes, BSR = burst spike rate, BD = Burst duration, BR = Burst rate, NBR = Network burst rate, NBD = Network burst duration, NBR = Network burst duration, IBI = inter-burst-interval, NIBI = Network burst IBI, $CV_{NIBI}$ = regularity of the network burst, RT = Rise Time, DT = decay time. DIV = Days in vitro.
